# Modeling of symbiotic bacterial biofilm growth with an example of the *Streptococcus-Veillonella* sp. system

**DOI:** 10.1101/2020.11.16.384172

**Authors:** Dianlei Feng, Insa Neuweiler, Regina Nogueira, Udo Nackenhorst

**Affiliations:** Institute of Fluid Mechanics and Environmental Physics in Civil Engineering, Leibniz Universität Hannover, Germany; Institute for Sanitary Engineering and Waste Management, Gottfried Wilhelm Leibniz Universität Hannover, Germany; Institute of Mechanics and Computational Mechanics, Leibniz Universität Hannover, Germany

## Abstract

We present a multi-dimensional continuum mathematical model for modeling the growth of a symbiotic biofilm system. We take a dual-species namely, the *Streptococcus - Veillonella* sp. biofilm system as an example for numerical investigations. The presented model describes both the cooperation and competition between these species of bacteria. The coupled partial differential equations are solved by using an integrative finite element numerical strategy. Numerical examples are carried out for studying the evolution and distribution of the bio-components. The results demonstrate that the presented model is capable of describing the symbiotic behavior of the biofilm system. However, homogenized numerical solutions are observed locally. To study the homogenization behavior of the model, numerical investigations regarding on how random initial biomass distribution influences the homogenization process are carried out. We found that a smaller correlation length of the initial biomass distribution leads to faster homogenization of the solution globally, however, shows more fluctuated biomass profiles along the biofilm thickness direction. More realistic scenarios with bacteria in patches are also investigated numerically in this study.

## 1 Introduction

More than 90% of microbes live in biofilms which can be defined as “assemblages of bacterial cells attached to a surface and enclosed in an adhesive matrix secreted by the cells”(Madigan, 2012). A biofilm *in natura* is usually found as a multi-component, multi-species, heterotopic matter with multi-phase properties. Competition and cooperation among species of bacteria are normally involved in multi-species biofilm systems (Yang et al., 2011). A biological system is symbiotic when cooperation happens between two different organisms.

Mathematical and numerical modeling is a powerful tool for understanding both the physical and bio-chemical processes during the formation and development of the biofilms. Many modeling strategies and models have been developed for describing biofilm processes. However, mathematical modeling of symbiotic biofilm systems has not been well studied. Modeling a symbiotic biofilm system naturally requires the consideration of a multi-species biofilm problem which is usually challenging, especially for multi-dimensional scenarios.

Several studies on modeling multi-dimensional multi-species biofilm system have been carried out in the literature. (Noguera et al., 1999) presented a Cellular Automaton (CA) model for modeling a dual-species *(D. vulgaris-M. formicicum)* biofilm system. Recently, (Tang and Liu, 2017) developed a multi-species multi-dimensional CA model to study syntrophic and dissimilatory metal reducing bacterial bioiflm system. In their study, a syntrophic ecological system is considered. (Martin et al., 2017) studied a dual-species *(S. gordonii-P. gingivalis)* oral biofilm system with a further developed CA model. Three models on the relationship between these two species of bacteria, namely independence for substrates, competition for substrate, and inhibition of one to another, are compared in their study. Individual-based Modeling (IbM) have also been used for studying various multi-species biofilm problems, such as modeling ammonium oxidizer bacteria (AOB) and nitrite oxidizer bacteria (NOB) system (Picioreanu et al., 2004), modeling dormant cell formation (Chihara et al., 2015) process, investigation of denitrifying bacteria – sulfate reducing bacteria – methanogens biofilm systems (Martin et al., 2015) and so forth. These models are not continuum and are usually categorized as the so-called discrete element based models (Fujikawa, 1994; Kreft et al., 2001; Noguera et al., 1999; Picioreanu et al., 1998; Wimpenny and Colasanti, 1997).

Although the reaction model presented in this paper can also be applied with discrete element based models, we present our model within the framework of continuum biofilm models which are fully described by partial differential equations (Alpkvist and Klapper, 2007; Cogan, 2004; Duddu et al., 2009; Eberl et al., 2001; Klapper and Dockery, 2002; Lindley et al., 2012; Wanner and Gujer, 1986; Zhang et al., 2008b). (Alpkvist and Klapper, 2007) presented a multi-dimensional multi-species biofilm model and applied it for studying an autotrophs-heterotrophs-inert bifilm system. The growth of biofilm is modeled as an advective movement of a potential flow due to the production or reduction of biomass. Instead of modeling the biofilms grow advectively, (Rahman et al., 2015) presented a multi-species multi-dimensional continuum diffusion-reaction model. A cross-diffusion process is additionally introduced into the model to overcome the internal over-mixing problem. The over-mixing of different species of biomass is a common flaw in many diffusion-reaction continuum models as well as some discrete element based models for multi-species multi-dimensional biofilm modeling (Tang and Valocchi, 2013). Since the advection-reaction biofilm model presented by Alpkvist and Klapper (Alpkvist and Klapper, 2007) does not involve physical diffusion, it is expected to be less diffusive, although the numerical dissipation is not avoidable. However, when the reactions become complex, whether locally homogenized biomass distribution will be obtained or not is still an open question. We look into such a problem by studying the presented symbiotic biofilm system with an advection-reaction type model, and one goal of this study is to investigate the solution behaviors of the model.

We take an oral dual-species, namely the *Streptococcus-Veillonella* sp. biofilm system (Chalmers et al., 2008), as an example of modeling symbiotic biofilm systems. Both *Streptococcus* sp. and *Veillonella* sp. play vital roles in the formation of dental biofilms. Dental bacterial biofilms are often of special interested due to their roles in the treatments of dental implants and diseases (Kommerein et al., 2017; Paquette et al., 2006). Different from many other types of biofilms, the bacteria that form the dental biofilms need salivary glycoproteins to attach to the teeth surface. Meanwhile, saliva is also the main nutrient source for the microbes in oral cavities (Chalmers, 2008;

Chalmers et al., 2008). Among the *Streptococcus* group bacteria, *Streptococcus gordonii* is a well-known Gram-Positive commensal bacterium in oral cavities which can cause caries and demineralization of teeth (Nascimento et al., 2009). *Streptococcus gordonii* is also the first batch of bacteria that attach to the teeth surface and is known as a species of early colonizers (Kolenbrander et al., 2010). The early colonizers can build up a foundation for the later colonizers to attach to the biofilm structure. This makes them important for studying oral biofilms.

Similar to the *S. gordonii, Veillonella* sp. is also a group of early colonizers which congregate with *S. gordonii* (Periasamy and Kolenbrander, 2010). It has been discovered that the *Veillonella* sp. cannot grow alone in saliva. However, it colonizes when there is *Strptococcus* sp. (e.g. *S. gordonii)* coexisting in the system (Chalmers et al., 2008). Similar behavior has also been observed in (Kara et al., 2007; Mashima and Nakazawa, 2015). A biological interpretation of such phenomenon is that the *Veillonella* sp. cannot utilize the carbon source in saliva directly but can ferment the metabolic production of *S. gordonii,* namely lactic acid, instead (Periasamy and Kolenbrander, 2010). It is worth noting that the lactic acid not only provides a carbon source for the growth of the *Veillonella* sp. but also causes a decrease of the pH in the system. It has been experimentally observed that the *Veillonella* sp. can tolerate more acid than the *S. gordonii* (Bradshaw and Marsh, 1998). The presence of the *Veillonella* which utilizes the lactic acid may thus be beneficial to the growth of the *S. gordonii* indirectly.

In this study, we develop a mathematical model for the symbiotic *S. gordonii – Veillonella* biofilm system under the hypothesis that the lactic acid has a negative influence on the growth of *S. gordonii* and thus the symbiotic biofilm system studied in this paper is essentially a mutualistic ecological system. The flow velocity in oral cavities is known to be very small and does not have much impact on the growth of the *S. gordonii* biofilm within a typical characteristic period (12h-24h) (Rath et al., 2017). The mathematical model developed by Alpkvist and Klapper (Alpkvist and Klapper, 2007) has been validated for the *S. gordonii* biofilm (Feng et al., 2018; Rath et al., 2017). Many studies have reported that the *Streptococcus* sp. dominates in a multi-species biofilm, which consists of the *Streptococcus* sp. and *Veillonella* sp. (Chalmers et al., 2008; Kommerein et al., 2017; Periasamy and Kolenbrander, 2010). For this reason, we develop the symbiotic biofilm model based on the advection-reaction biofilm growth model.

The model presented in this paper can be used for quantitatively understanding of the *Streptococcus-Veillonella* symbiotic biofillm system and in general can be modified to model a variety of biofilm systems with similar bio-chemical properties. To the best of our knowledge, a continuum model for such kind of symbiotic biofilm systems has not been well developed. This study may shed light on understanding the numerical behaviors of multi-dimensional multi-species continuum (advection-reaction) biofilm models which involve complex reactions.

This paper is structured as follows. Details of the model will be presented in section 2. Numerical results are presented and investigated in section 3. The symbiotic biofilm model is highly non-linear. We observed a homogenized numerical solution of biomass at a later time. To better understand the solution behaviors, we investigated how the initial condition (see section 3.2) and the symbiotic reaction relationship (see section 3.4) influence the homogenization of the solutions. In section 4, the results are summarized and discussed. Further detailed information regarding the dimensionless form of the governing equations, as well as the corresponding numerical method, are presented in Appendix A and B.

## 2 Mathematical model

### 2.1 Model assumptions

The *Streptococcus-Veillonella* biofilm system is illustrated in Figure 2. Two biomass components, namely the *S. gordonii* and *V. PK1910* interact with each other in a mutualistic relationship. Especially, *S. gordonii* produces lactic acid, which plays a vital role in the chain of the symbiotic bio-chemical process is modeled explicitly. Despite the saliva cannot function as a carbon source for the *Veillonella* sp., saliva still may provide other nutrients for the growth of both species due to its complex mixture substance property. Therefore, a competition of the saliva between *S. gordonii* and *Veillonella* sp. is considered in the model. On the other hand, *Veillonella* sp. helps to prevent the environment from being too acid, and thus benefits the growth of *S. gordonii*. We develop the mathematical model based on the following assumptions:

- A layer of biomass with a finite thickness has been presented as an initial condition. The model does not capture the process in which individual bacteria grow into a biofilm. On the other hand, the growth process after forming an initial layer of biofilm is modeled.
- Two species of bacteria, namely the *Streptococcus* sp., and *Veillonella* sp. are considered in the model. The volume fractions of the bacteria sum to one inside of the biofilm, which implies that the pore space and EPS are not explicitly modeled (Alpkvist and Klapper, 2007).
- Saliva is considered as a common nutrient source consumed by both species of bacteria. However, the growth of *Veillonella* is limited by both the nutrient and the lactic acid, which is produced by *S’treptoccus* sp (Chalmers et al., 2008).
- The concentration of hydrogen is not modeled explicitly, but the concentration of the lactic acid is considered instead as an indicator for it (Gordeeva et al., 2017).
- The fluid-structure interaction process is not considered. This assumption is based on our previous experimental observations of the formation process of *S. gordonii* (Rath et al., 2017).
- No new void spaces of bacteria can develop during the growth of the biofilm. In other words, the biofilm domain is simply connected (see Figure 1).
- The nutrients in saliva are fully mixed with a distance above the biofilm. A Dirichlet boundary condition of the nutrients thus exists (Picioreanu et al., 2006).
- The growth velocity is irrotational (Klapper and Dockery, 2002), and different species of bacteria have the same advection velocity (Alpkvist and Klapper, 2007).
- Diffusion coefficients are constant both inside and outside of the biofilm (Alpkvist and Klapper, 2007).

**Figure 1.**
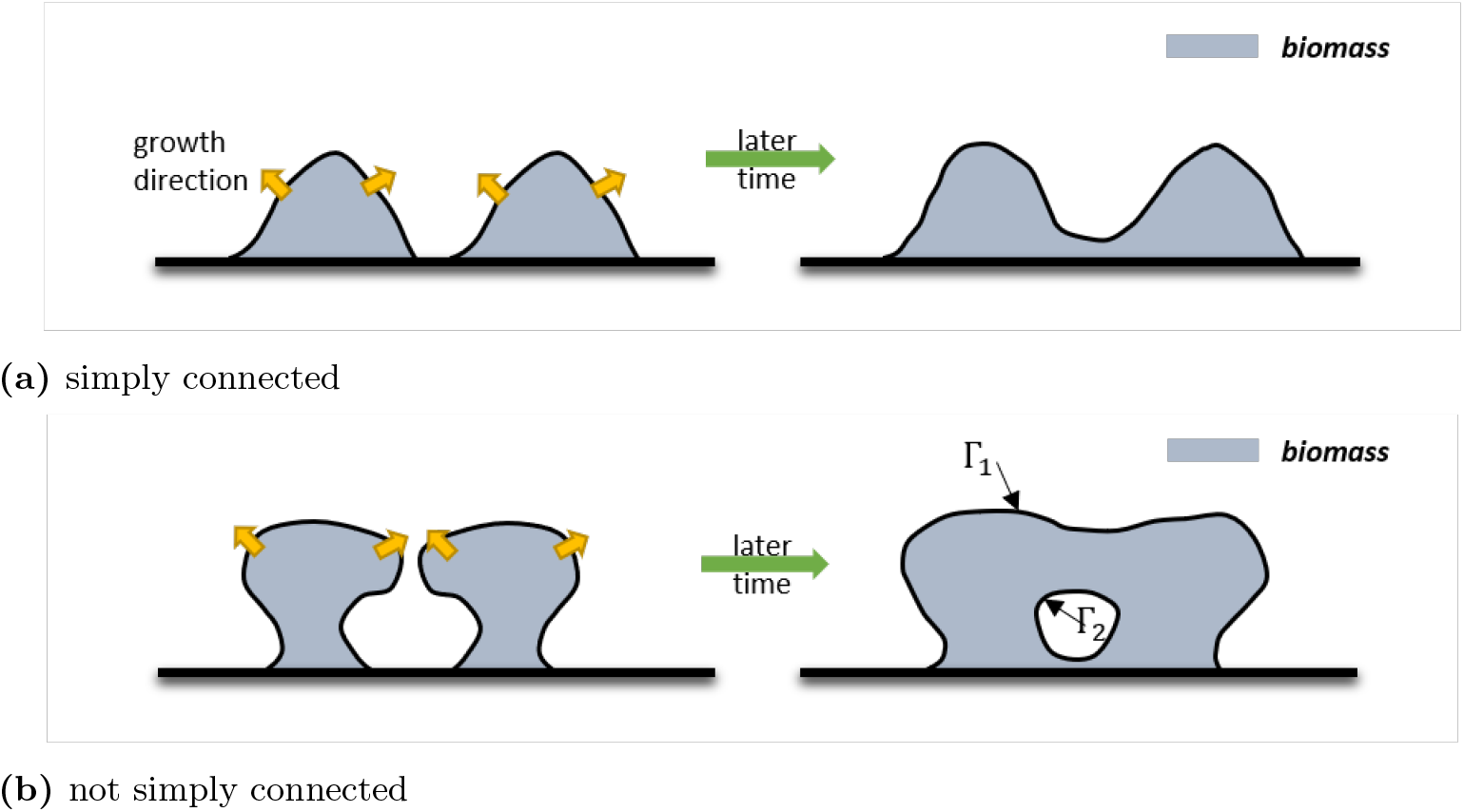
Typical biofilm patterns generated during the merge of two colonies.

**Figure 2.**
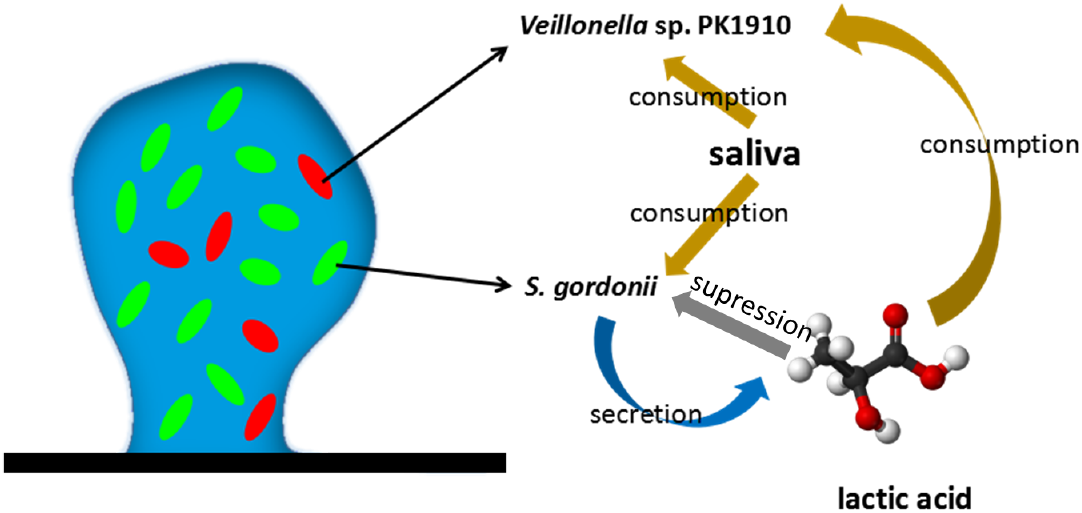
Illustration of the symbiotic biofilm system. The *Veillonella* sp. PK1910 and *Streptococcus gordonii* are taken as example species of the bacteria in the illustration. Both species of bacteria compete for the nutrients in saliva. The *Streptococcus gordonii* produces lactic acid, which is a carbon source for *Veillonella* sp. The lactic acid also prevents the growth of *gordonii.*

To make the model simple, we introduced assumptions more than necessary. One can include the EPS as an additional biomass component in the model, as in (Xavier et al., 2005). It is also convenient to assign heterogeneous modeling parameters, such as diffusion coefficient, density, or even growth rate in the model. The pH can also be modeled by slightly extending the model, as presented in (Khassehkhan and Eberl, 2008).

The main limitation of the presented model is the requirement of being simply connected (biofilm domain) during biofilm growth. This is mostly due to the difficulty of enforcing proper potential (or pressure) boundary conditions for the newly generated biofilm-fluid surface. Such a problem may come up when trying to model the merge of different colonies. As illustrated in Figure 1b, when there is a new biofilm-fluid interface (Γ_2_) generated during the merge of colonies, it is not straightforward to enforce boundary conditions of the potential on Γ_1_ and Γ_2_. On the other hand, the model can be used to model the merge of different colonies if only the scenario, as shown in Figure 1a, occurs. Modeling the merge of colonies is beyond the aim of this paper. For the readers interested in this topic, we refer to the three-dimension numerical study in (Alpkvist and Klapper, 2007).

### 2.2 Model domain

To build up the model based on the above assumptions, the mass balance of each substrate and component of biomass in the biofilm is considered. Eventually, the mathematical model, which is essentially a free boundary problem, consists of five partial differential equations (PDEs) including four coupled nonlinear time dependent advection-diffusion-reaction (or advection-reaction) equations and a Poisson’s equation.

Taking a 2D model as an example, the biofilm is modeled within a computational domain of Ω = {**x** = (*x, z*): 0 ≤ *x* ≤ *W*, 0 ≤ *z* ≤ *H*} with the corresponding boundary *∂*Ω. The top border is denoted as Γ_*ht*_ and the other sides are denoted by 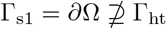. The outward-pointing normal vectors of Γ_s1_ and *∂*Ω are denoted as **n**_*s*1_ and **n**_*s*2_ respectively. A sub-domain B_t_ ∈ Ω refers to the biofilm domain above which a bulk fluid domain F_t_ is assumed. The biofilm-fluid interface Γ_int_ = B_t_ ⋂ F_t_, which is a free boundary, moves over time induced by the growth of the biofilm.

### 2.3 Mass balance of biomass

The growth of both *S. gordonii* and *Veillonella* are modeled as advective movements driven by the production and reduction of biomass inside of the biofilm. Writing the mass balance equations of *S. gordonii* and *Veillonella* sp. in terms of their volume fraction *ϑ* reads

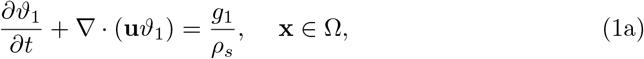

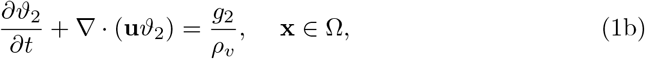

where **u** [*LT*^-1^] denotes the growth velocity of the biofilm, indexes “1” and “2” are used to refer physical quantities of the *S. gordonii* and *Veillonella* bacteria respectively and, *ρ_s_* and *ρ_v_* denote the density of the corresponding biomass. In equation (1), *g* refers to the mass production/reduction rate of the biomass. A no-flow boundary is applied for both equation (1a) and equation (1b)

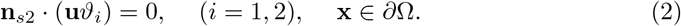

It has been experimentally found that the *S. gordonii* can tolerate less acid than the *Veillonella* sp. and the *Veillonella* sp. may become dominant even in the environment with pH < 5.0 (Bradshaw and Marsh, 1998). The growth of *S. gordonii* is sensitive to the acid and, on the other hand, *Veillonella* can grow in a small pH environment. Therefore, we only model the negative influence of lactic acid on the growth of *S. gordonii.* The mass production process of *S. gordonii* is limited by the concentration of saliva (*s*_1_) and suppressed by the lactic acid (*s*_2_)

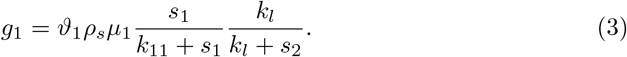

In equation (3), *μ*_1_ [*T*^-1^] and *k*_11_ [*ML*^-3^] refer to the maximum growth rate and the half-rate constant of the *S. gordonii* in the Monod kinetic respectively. We introduce a parameter *k_l_* [*ML*^-3^] for modeling the suppression process induced by the lactic acid. A benefit of choosing *g*_1_ in the form as equation (3) is, when *s*_2_ → +∞, *g*_1_ → 0. This means the growth of the *S. gordonii* stops when the concentration of the lactic acid is infinitely large.

The growth of *Veillonella* is limited by both the concentration of the nutrients in saliva *s*_1_ and the concentration of the lactic acid *s*_2_

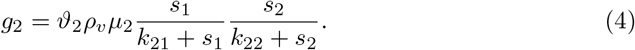

In equation (4), *μ*_2_ [*T*^1^] denotes the maximum growth rate of *Veillonella* and *k*_21_ [*ML*^-3^] and *k*_22_ [*ML*^-3^] are the half-rate constant in the Monod kinetic.

### 2.4 Mass balance of saliva and lactic acid

The chemical reactions of components in the saliva environment are very complex, and the corresponding reaction parameters are mostly unknown. Instead of describing detailed chemical reactions of all involved chemical components in saliva, we consider the saliva itself as one substance demanded by both species of bacteria. Since the time scale of the transport process of the substrate (solutions) is much smaller than the time scale of the biofilm growth (Picioreanu et al., 2000), the mass balance of the saliva is modeled as a stationary diffusion-reaction process as

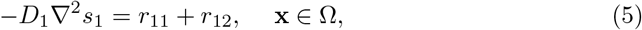

where *r*_11_ and *r*_12_ [*ML*^-3^*T*^-1^] denote the saliva (nutrients in saliva) consumption rates by *S. gordonii* and *Veillonella* sp. respectively, and *D*_1_ [*L*^2^*T*^-1^] denotes the diffusion coefficient of the saliva. As a remark, one can also write (5) as a time-dependent equation (Wanner and Reichert, 1996).

The nutrient in saliva comes from the outside of the modeling domain Ω. A constant concentration 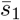 is set at the top of the computational domain Γ_*ht*_ (as shown in Figure 3)

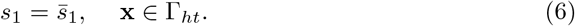

**Figure 3.**
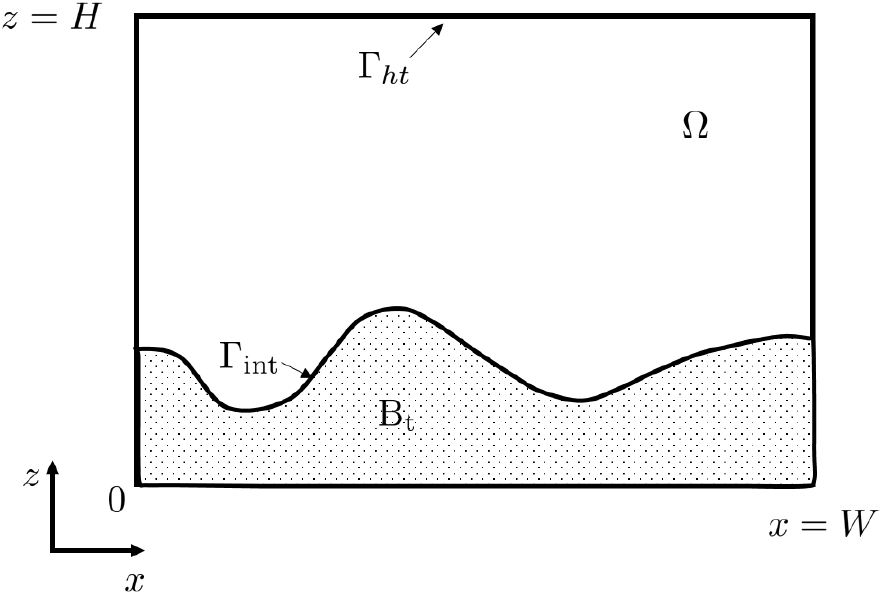
Two dimensional illustration of the computational domains of the symbiotic biofilm model.

A no-flow boundary is applied on the other borders of the computational domain as

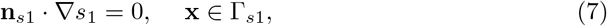

where **n**_*s*1_ denotes the normal vector of Γ_*s*1_.

The reaction term corresponding to the consumption of saliva by *S. gordonii* is described by using the Monod kinetics and reads

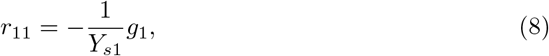

where *Y*_*s*1_[—] denotes the corresponding yield of the *S. gordonii* by consuming rate of the saliva. Similarly, the consumption rate of the saliva due to the *Veillonella* reads

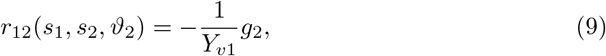

where *Y*_*v*1_ [—] is the yield of the *Veillonella.*

The lactic acid which is secreted by the *S. gordonii* accumulates over time in the system. If a batch system is considered, the lactic acid is produced inside of the biofilm domain B_t_ and stays in the system. In this case, the mass balance of the lactic acid cannot be modeled as a quasi-steady state since the absence of the Dirichlet boundary condition. For these reasons, the mass balance of the acid has to be modeled as a time dependent process. The mass transport of acid is diffusion-reaction dominated.Therefore, one can simplify the model by neglecting the advection process and achieve comparable, accurate results. Moreover, the production and consumption of the lactic acid by *S. gordonii* and *Veillonella* sp. are also considered respectively. The mass balance of the lactic acid reads

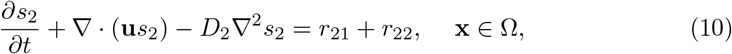

where *D*_2_ [*L*^2^*T*^-1^] is the diffusion coefficient of the lactic acid. *r*_21_ [*ML*^-3^*T*^-1^] and *r*_22_ [*ML*^-3^*T*^-1^] describe the production of the lactic acid by *S. gordonii* and the consumption rate of the acid by the *Veillonella* respectively.

*S. gordonii* grows by consuming saliva and producing lactic acid at the same time. Thus the reaction term *r*_21_ can be written as

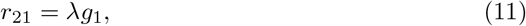

where λ [—] is a parameter which describes the mass production rate of acid by unit mass of *S. gordonii.* Only the *Veillonella* sp. in the system consumes lactic acid, therefore, the reaction kinetic can be written as

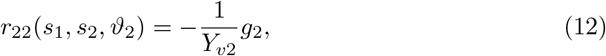

where *Y*_*v*2_ refers to the yield of *Veillonella* by consuming the lactic acid. A no-flow boundary is applied on *∂*Ω for equation (10) as

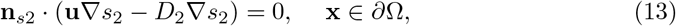

where **n**_*s*2_ denotes the normal vector on *∂*Ω.

We applied a no-flow boundary condition in the saliva’s mass balance equation. Such a boundary condition implies that a batch system is modeled. For flow chamber systems, one may also invoke a Dirichlet boundary for the acid (see later discussions in section 3.1.2). Alternatively, one can apply a diffusive boundary layer (Feng et al., 2017) which essentially is a Robin boundary condition (Ghasemi et al., 2018; Wanner and Gujer, 1986) for allowing the acid to leave the domain.

### 2.5 Potential equation for the movement of the biofilm

We model the movement of the biofilm as a potential flow driven by the production of the biomass inside of the biofilm following (Alpkvist and Klapper, 2007; Klapper and Dockery, 2002). It is assumed that the two species of bacteria compose the whole biofilm. Therefore, the volume fractions of the *S. gordonii* and *Veillonella* sum up to one in the biofilm domain B_t_ as

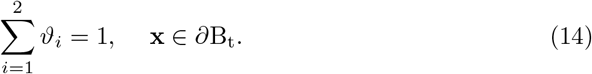

Following the previous potential flow assumption, the biofilm growth velocity is described as the gradient of the potential Φ. Substituting equations (1a) and (1b) into equation (14) yields the potential equation

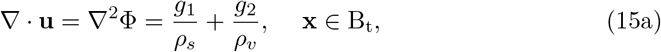

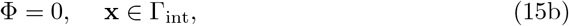

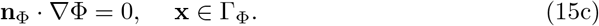

In equation (15), **n**_Φ_ denotes the norm vector of **Γ**_Φ_. Additionally, **u** = 0 in the domain outside of the biofilm B_t_.

### 2.6 Dimensionless form of the governing equations

Equations (1), (5), (10) and (15a) together with their corresponding boundary conditions compose the full mathematical model which describes the symbiotic biofilm system illustrated in Figure 2.

To obtain the dimensionless form of the governing equations, we introduce the following dimensionless variables

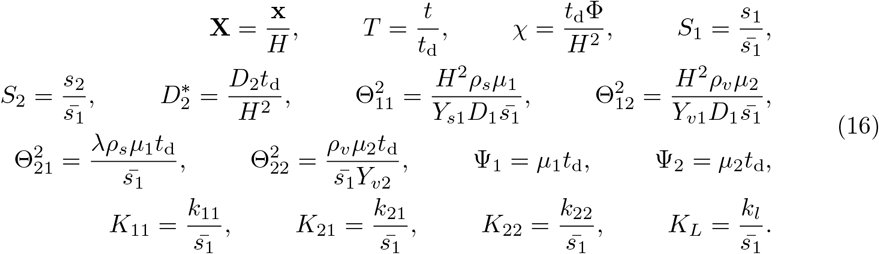

In equation (16), *H* and *t*_d_ are characteristic length and time which are taken as the height of the computational domain and the time of a day respectively in this study.

Substituting equation (16) into equations (1), (5), (10) and (15a) yields the dimensionless form of the governing equations. The detailed dimensionless governing equations are listed in Appendix A, and the corresponding numerical strategy for solving these dimensionless equations is presented in Appendix B and C.

## 3 Results

We investigate the solution behaviors of the mathematical model in this section. An initially evenly mixed scenario has been investigated in section 3.1. It turns out that the solution is spatially homogenized at a later time. To better understand the solution’s homogenization property, we studied how different random initial conditions influence the solutions in section 3.2. We also look into more realistic scenarios with an initial condition of one species embedded in another and investigate the role of symbiotic reactions in the generation of homogenized solutions (in section 3.3 and 3.4).

Two-dimensional simulations are carried out with the modeling parameters listed in Table 1. For all numerical simulations, a 200 × 200 spatial mesh with four-node bilinear elements is applied and the time step size is taken as 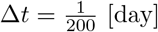. The TDG-FIC scheme (see Appendix B) with a 3rd order time accuracy is applied for solving the governing equations. In this study, we aim at investigating the reactive dynamics of the symbiotic biofilm system instead of a specific experimental scenario. We apply an initial condition for *S*_2_ by assuming no lactic acid in the system as

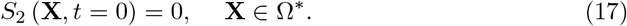

**Table 1.** Parameters used for two-dimensional symbiotic biofilm simulation

| Quantity name | Symbol | Value | Unit | Source |
| --- | --- | --- | --- | --- |
| Length of the computational domain | $W$ | 300 | $\mu\text{m}$ | (Alpkvist and Klap-<br>per, 2007) |
| Height of the computational domain | $H$ | 300 | $\mu\text{m}$ | (Alpkvist and Klap-<br>per, 2007) |
| Saliva concentration at the top boundary | $\bar{s}_1$ | 10 | $\text{kg}/\text{m}^3$ | estimated |
| Diffusion coefficient of saliva | $D_1$ | $5 \times 10^{-10}$ | $\text{m}^2/\text{s}$ | (Corbin et al., 2011) |
| Diffusion coefficient of lactic acid | $D_2$ | $1 \times 10^{-9}$ | $\text{m}^2/\text{s}$ | (Ribeiro et al., 2005) |
| Density of *S. gordonii* | $\rho_s$ | 1100 | $\text{kg}/\text{m}^3$ | estimated from<br>(Duddu et al., 2009) |
| Density of *Veillonella* | $\rho_v$ | 1100 | $\text{kg}/\text{m}^3$ | estimated from<br>(Duddu et al., 2009) |
| Biofilm yield of *S. gordonii* by consuming saliva | $Y_{s_1}$ | $10^{-1}$ | $[-]$ | estimated |
| Biofilm yield of *Veillonella* by consuming saliva | $Y_{v_1}$ | $10^{-1}$ | $[-]$ | estimated |
| Biofilm yield of *Veillonella* by consuming lactic acid | $Y_{v_2}$ | $10^{-1}$ | $[-]$ | estimated |
| Production coefficient of lactic acid | $\lambda$ | $5 \times 10^{-1}$ | $[-]$ | estimated |
| Maximum growth rate of *S. gordonii* | $\mu_1$ | $3 \times 10^{-5}$ | $\text{s}^{-1}$ | (Rath et al., 2017) |
| Maximum growth rate of *Veillonella* | $\mu_2$ | $8 \times 10^{-5}$ | $\text{s}^{-1}$ | estimated |
| Monod half-rate constant | $k_{11}$ | 0.878 | $\text{kg}/\text{m}^3$ | (Rath et al., 2017) |
| Monod half-rate constant | $k_{21}$ | 0.878 | $\text{kg}/\text{m}^3$ | estimated |
| Monod half-rate constant | $k_{22}$ | 0.0878 | $\text{kg}/\text{m}^3$ | estimated |
| Monod half-rate constant | $k_l$ | 1 | $\text{kg}/\text{m}^3$ | estimated |

### 3.1 Evenly mixed case

A wave shape initial biofilm-fluid interface in the dimensionless domain is applied

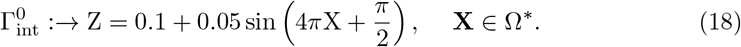

We also consider that these two species of bacteria are initially evenly mixed as

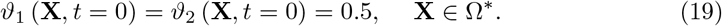

#### 3.1.1 Biomass evolution and distribution

Simulation results of biomass distribution of both *S. gordonii* and *Veillonella* at early and late times are shown in Figure 4. Since the numerical solutions are symmetric regarding the middle line *X* = 0.5, only the half plane of a solution of each species of biomass is shown in the figure. The white curve in Figure 4 represents the biofilm-fluid interface. The simulation results demonstrate that a flat biofilm pattern is generated after the growth of the biofilm for a certain time. This is due to the nutrients are not scarce in the system. However, we still can observe that the waved biofilm-fluid interface has quite an impact on the local distribution of the lactic acid (see later discussions).

**Figure 4.**
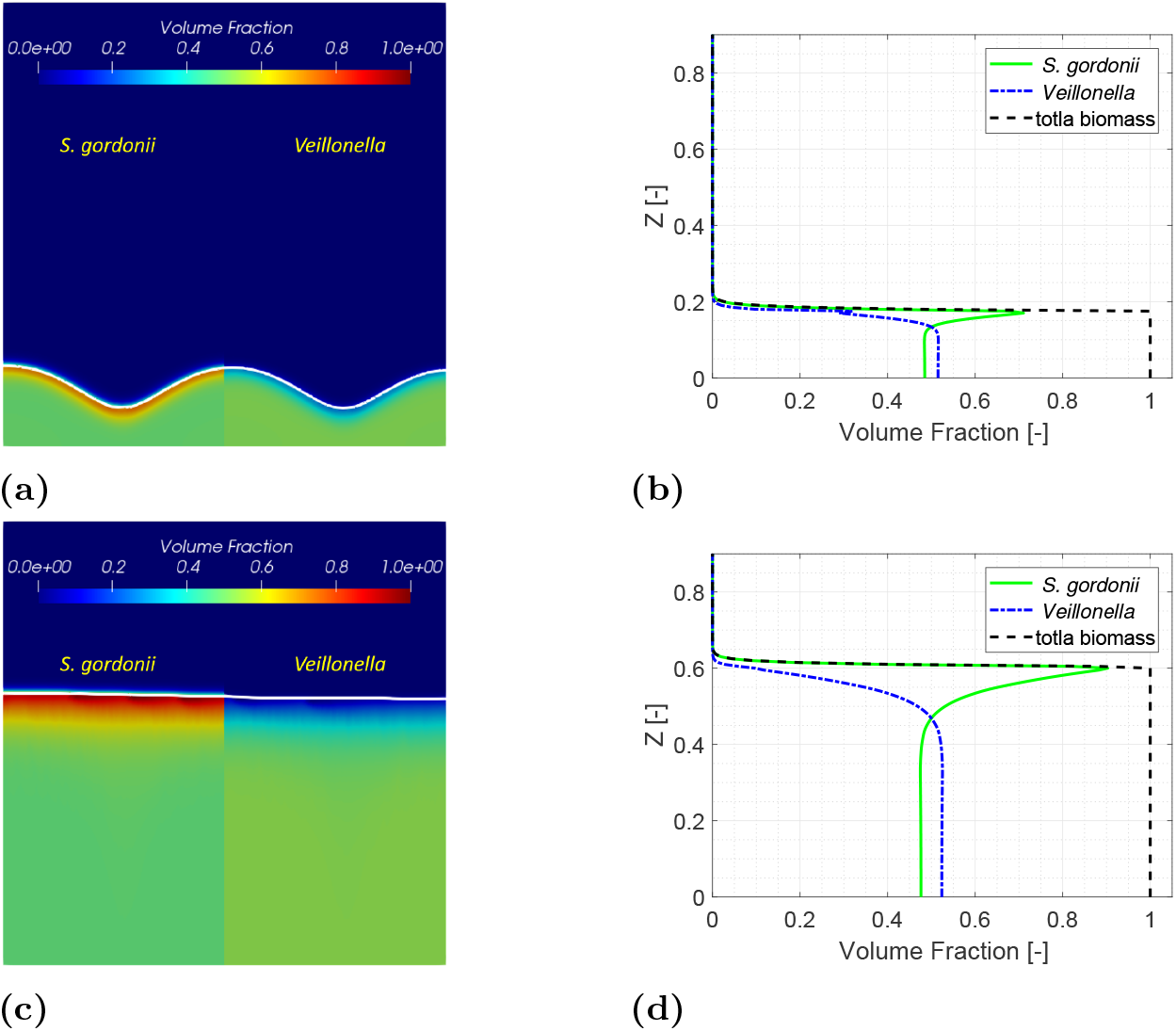
Simulation results of the symbiotic biofilm system at different times (from top to bottom are at *t* = 9.6h(*T* = 0.4) and *t* = 48h(*T* = 2.0)). Left column: Volume fraction distributions of *S. gordonii* and *Veillonella;* Right column: Volume fraction profiles of *S. gordonii* and *Veillonella* and total biomass in depth at *X* = 0.5.

Biomass volume fraction profiles of *S. gordonii*, *Veillonella* and total biomass at different times along *X* = 0.5 are shown in the right column in Figure 4. It is observed that *S. gordonii* moves up as a result of winning the competition for nutrients in saliva in the vicinity of the biofilm-fluid interface. Such a phenomenon has also been observed in experimental studies (Chalmers, 2008) as shown in Figure 5. In the figure, *S. gordonii* and *Veillonella* PK1910 are colored in green and blue respectively. We observe that the *S. gordonii* locates above the *Veillonella* in the areas within the red squares in Figure 5. As a remark, we are not trying to reproduce the biofilm pattern shown in Figure 5 in this paper. The pattern of biofilms highly depends on nutrients conditions.

**Figure 5.**
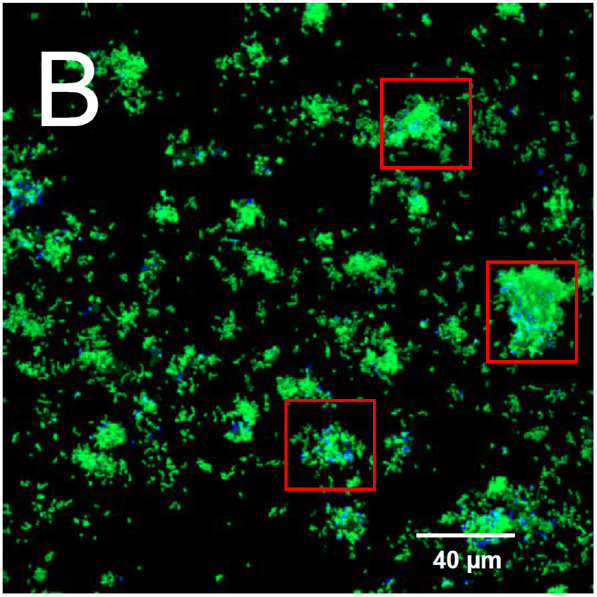
Spatial distribution of *S. gordonii* (in green) and *Veillonella* PK1910 (in blue) observed with confocal laser scanning microscopy in 2h flowcell biofilms (Chalmers, 2008).

A diluted saliva (Chalmers, 2008) is used as a nutrient in the experiments corresponding to Figure 5. We applied the calibrated model parameters corresponding to the growth of *S. gordonii* from our previous experimental study(Rath et al., 2017). The *S. gordonii* is cultured in the Tryptic Soy Broth medium instead of the diluted saliva. Under such a condition, we find the *S. gordonii* build up a biofilm of layering pattern (Kommerein et al., 2017; Rath et al., 2017). It has been observed that the *Streptococcus* sp. dominates when they are coaggregated with the *Veillonella* sp. Therefore, all numerical examples in this paper are set up with layered initial conditions.

The inhibition parameter *k_l_* has a significant influence on the growth of both species of bacteria. According to equation (3), the lactic acid does not prevent the growth of *S. gordonii* when *k_l_* is an infinitely large number. In this case, one would expect the *S. gordonii* grows at a large rate. As shown in Figure 6, we observe a faster growth of both species of bacteria by using a larger value of *k_l_*. As a remark, it is not surprised to see that the *Veillonella* sp. also grows faster associatively with the *S. gordonii.* This is due to the mutualistic relationship between these two species.

**Figure 6.**
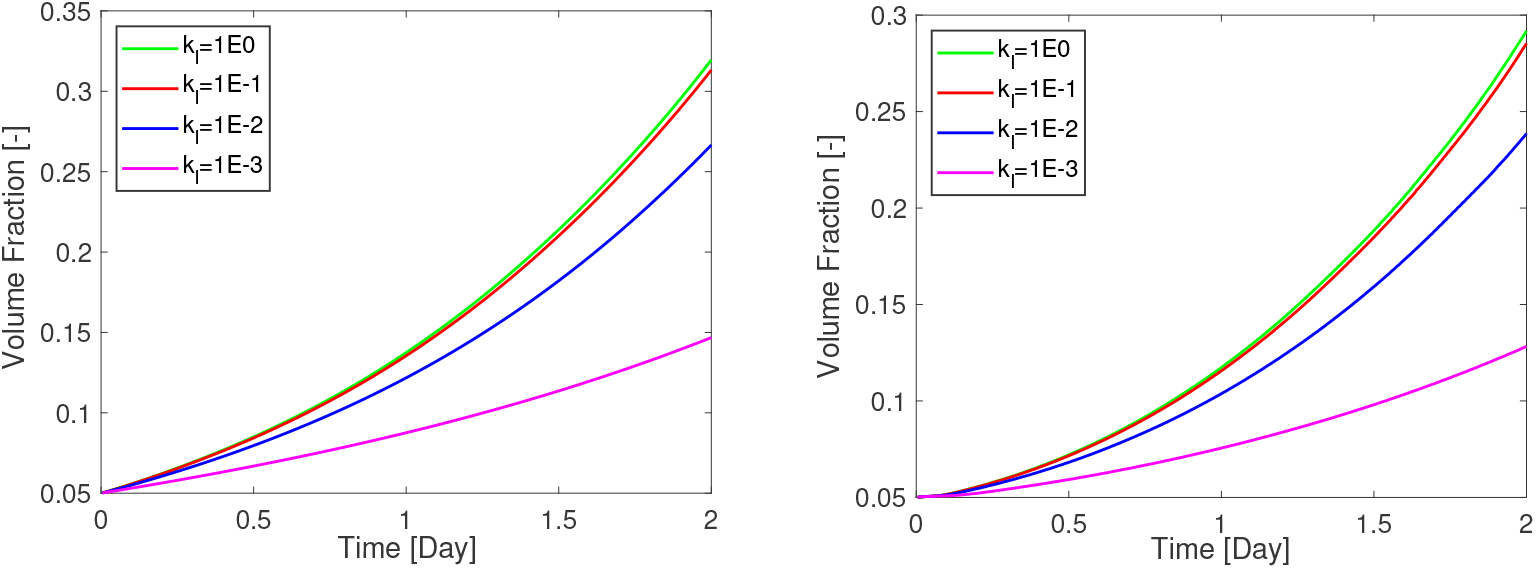
Sensitivity analysis of the inhibition parameter *k_l_* on the growth of bacteria.

It should be noted that the simulation results correspond to an evenly mixed initial biomass volume condition. In nature, it is observed that *S.gordonii* usually attaches to a surface first and *Veillonella* attaches to *Streptococcus* afterwards. They form the so-called “corn cob” pattern (Kolenbrander et al., 2010) *in vitro.* Such a pattern demonstrates that the initial condition of either the geometry of the biofilm-fluid interface or the biomass distribution in such a symbiotic biofilm system is heterogeneous. To study such complex heterogeneity properties that are influenced by the initial conditions, an attachment process with considerations of bacteria receptors has to be considered. The attachment process normally happens at an individual bacterium scale and can hardly be considered only with a continuum model. To capture the attachment process, modeling the whole biofilm system with a multi-scale model will be promising. However, this is beyond the scope of this paper.

#### 3.1.2 Lactic acid

Distribution of the lactic acid in terms of its dimensionless concentration *S*_2_ at different times is presented in Figure 7. The lactic acid is produced by the *S. gordonii* and consumed by the *Veillonella*. The acid is transported diffusively and accumulates in the fluid domain. It is observed that a higher concentration of acid accumulates above the biofilm-fluid interface, especially, after the *S. gordonii* becomes dominant at the top layer of the biofilm. Since there is more *S. gordonii* at the valley of the biofilm surface at *T* = 1.0 and *T* = 1.6, two punctiform sources of the acid at the biofilm valley are observed. This is why we observe the maximum acid concentration appears in the middle of the computational domain, and the acid diffuses to the top boundary. The *Veillonella* wins the competition at the lower part of the biofilm (see biomass profiles in Figure 4) where the lactic acid is largely consumed by the *Veillonella.* What should be noted is that the distribution of the lactic acid is also influenced by the biofilm patterns and the generation of biofilm patterns might be influenced by the distribution of the lactic acid as well.

**Figure 7.**
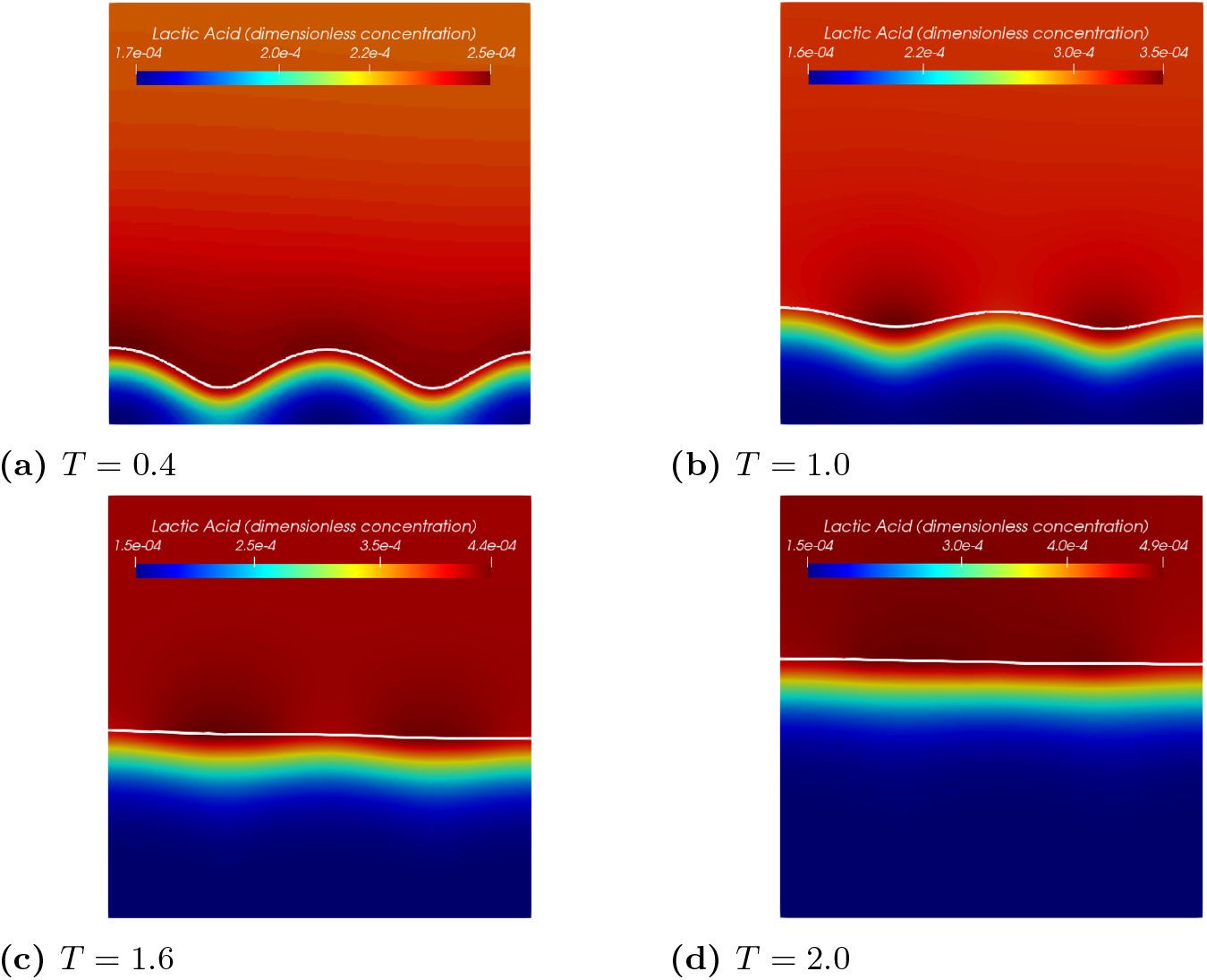
Distribution of lactic acid of the symbiotic biofilm system at different times (*T* = 0.4, 1.0, 1.6, 2.0). The white curve in the figures denotes the biofilm-fluid interface.

Figure 8 shows profiles of the lactic acid at *X* = 0.5 at different times. For each plot, the concentration of the lactic acid increases from the bottom of the biofilm to the top of it and decreases slightly afterwards in the bulk fluid. The profiles demonstrate that the location of the maximum concentration moves together with the biofilm interface. The profiles also show that the lactic acid accumulates in the bulk fluid domain over time. Comparing the two profiles of *t* = 1.6 day(*T* = 1.6) and *t* = 2.0 day(*T* = 2.0), the difference of the acid concentrations at the bottom of the biofilm is rather small. It seems the solution reaches a steady state locally there.

**Figure 8.**
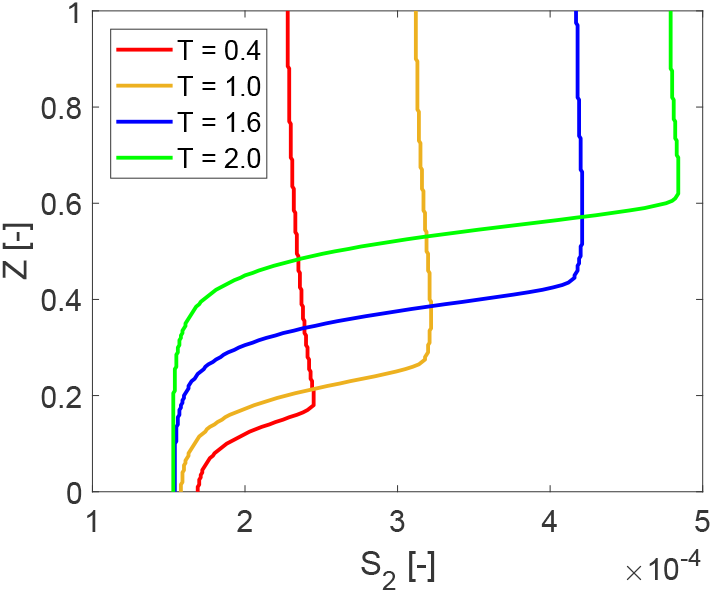
Dimensionless concentration profiles of lactic acid at *X* = 0.5 at different times (*T* = 0.4, 1.0, 1.6, 2.0) with a no-flow boundary condition.

Since we apply a no-flow boundary to model the acid transport in a batch system. The acid accumulates in the fluid domain over time. One can also apply a Dirichlet boundary for *s*_2_. By assuming the acid is removed immediately at the top of the computation domain, equation (13) can be replaced by

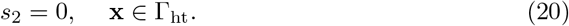

As shown in Figure 9, the lactic acid profiles are different than the profiles shown in Figure 8 in the fluid domain. We further compare the solutions of the lactic acid by using different boundary conditions in Figure 10. It turns out that different boundary conditions do not affect the acid profiles much in the biofilm.

**Figure 9.**
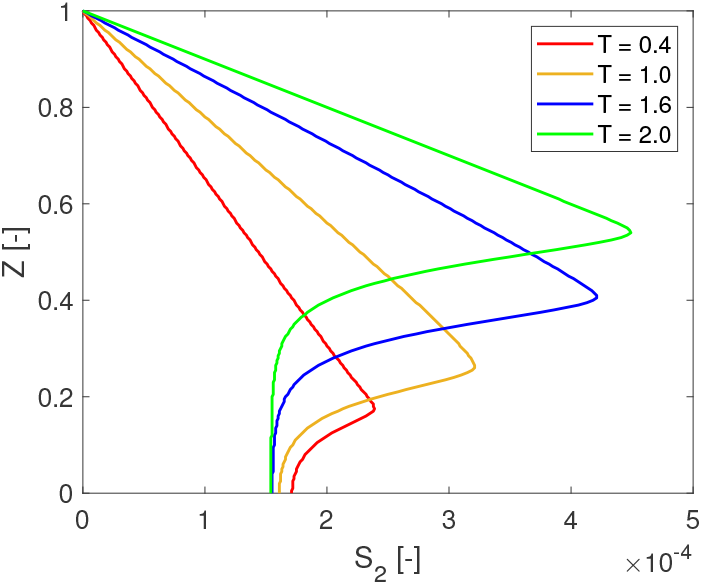
Dimensionless concentration profiles of lactic acid at *X* = 0.5 at different times (*T* = 0.4, 1.0, 1.6, 2.0) with a Dirichlet boundary condition.

**Figure 10.**
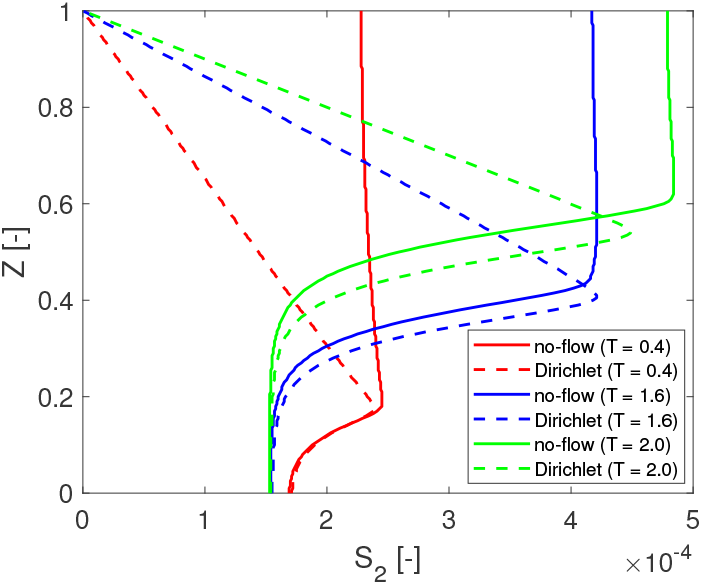
Comparison of lactic acid profiles at *X* = 0.5 with different boundary conditions at *T* = 0.4, 1.6 and 2.0.

The lactic acid plays a twofold role in biofilm growth. On the one hand, it is the carbon source for *Veillonella.* On the other hand, the acid inhibits the growth of *Streptococcus.* In this study, we apply a relatively large value of *k_l_* (see Table 1), which reduces the second effect. Since the acid does not accumulate in the bulk fluid when the Dirichlet boundary is applied, its concentration is relatively smaller, especially at a later time (e.g., the peak value of *S*_2_ at *T* =2.0). Therefore, the biofilm grows slower. The dimensionless thickness of the biofilm reduces 3% at *T* = 1.6 by using the Dirichlet boundary. At *T* =2.0, the reduction is increased to 6%. This study aims to investigate the solution behaviors, the boundary condition of the acid will not affect our investigation results.

The simulation results of this 2D example shed a light on the properties of the multi-dimensional numerical solutions of the presented biofilm model. First, heterogeneous biomass distribution is obtained with this model even though a fully mixed initial condition is applied. On the other hand, the biomass is homogenized locally in the biofilm. Second, the biofilm-fluid interface is flatted in time. However, the non-flat interface has an influence on the lactic acid distribution. Based on those observations, we ask ourselves: 1) Will the local homogenization process always happen in this model regardless of the initial biomass distribution? If so, will the initial distribution influence the homogenization process? 2) How will a heterogeneous initial biomass distribution influence the morphology of the biofilm interface as well as the components (including the biomass and the lactic acid) distributions during the growth?

### 3.2 Sensitivity to initial biomass distribution

In order to answer these questions raised in the previous section, we study the sensitivity of the presented mathematical model to different types of initial biomass distributions. As a matter of fact, the initial distribution of different species of bacteria in a biofilm can be never evenly mixed in nature. As shown in Figure 11, we consider three cases of the random initial distribution of biomass. Random fields are generated by using given different correlation lengths. The correlation length measures the distance of two correlated points (e.g., the points with correlated material properties or physical quantities). Thus, the correlation length is often seen as a measure of the roughness of surfaces. In this study, different correlation lengths of 2Δ*x*, 4Δ*x*, and 8Δ*x* are applied in each case, in which 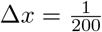 denotes the length of a spatial element in the horizontal direction. As a remark, the total initial mass (or volume) of each species of bacteria is the same. To simplify the problem, the initial biofilm-fluid interface is modeled as a flat line, and a constant initial biofilm thickness of 60 *μ*m (*Z* = 0.2) is assumed. Following our previous numerical setup, we assume that there is no lactic acid in the system initially.

**Figure 11.**
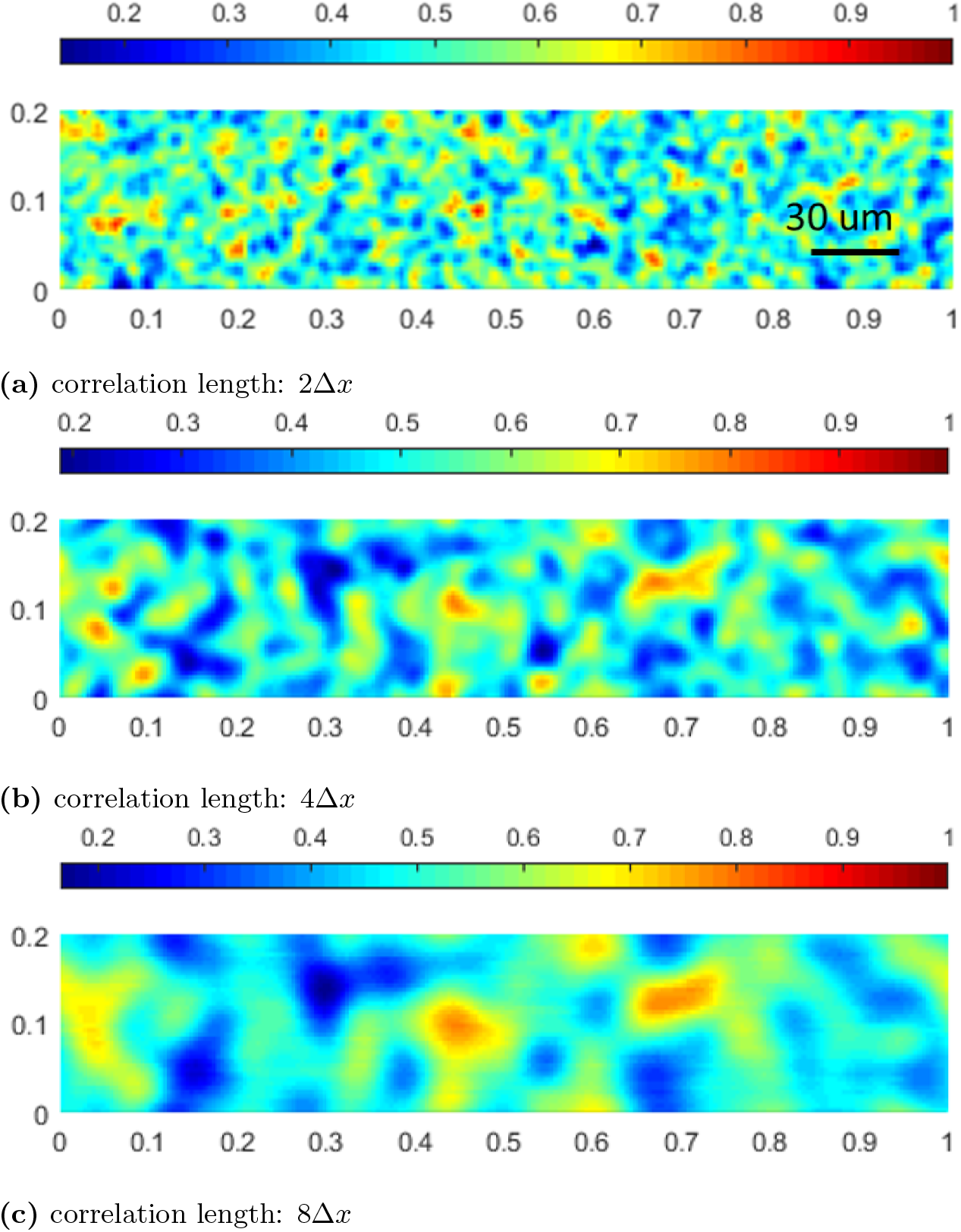
Initial random volume fraction distribution of *Veillonella* with different correlation lengths.

As shown in Figure 12, the *Veillonella* patches are stretched and thus generating plume-like structures during the biofilm formation in all three cases. This indicates that processes happen mainly in the vertical direction, and the horizontal movement is not obvious. By increasing the correlation length of the initial biomass distribution, the size of the plumes increases. This results in that the biomass volume fraction profiles along a vertical line, for instance at *x* = 150 *μ*m (*X* = 0.5) as shown in Figure 13, get less fluctuated with a larger correlation length. However, this does not mean that with a larger correlation length, the biomass homogenizes faster. In order to study how different biomass initial distributions influence the homogenization time, we compare the numerical solutions corresponding to the random initial distributions to the solution with an evenly mixed initial biomass distribution. Since the simulation involves moving domains, measuring the homogenization process is nontrivial. We calculate the difference of the volume fractions of the *Veillonella* between the evenly mixed and randomly distributed cases to measure the homogenization process

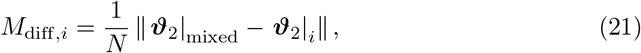

where *ϑ*_2_ is the solution vector of the volume fraction of *Veillonella, N* is the number of the unknowns in the vector and indexes “mixed” refers to the evenly mixed initial condition. The index *i* = 1, 2, 3 refers to scenarios with different correlation lengths of 2Δ*x*, 4Δ*x* and 8Δ*x*. Since the evenly mixed case always presents a locally homogenized solution which produces a comparable biofilm thickness evolution as the other cases at an early time, a smaller *M*_diff_ corresponds to a solution which is more homogenized.

**Figure 12.**
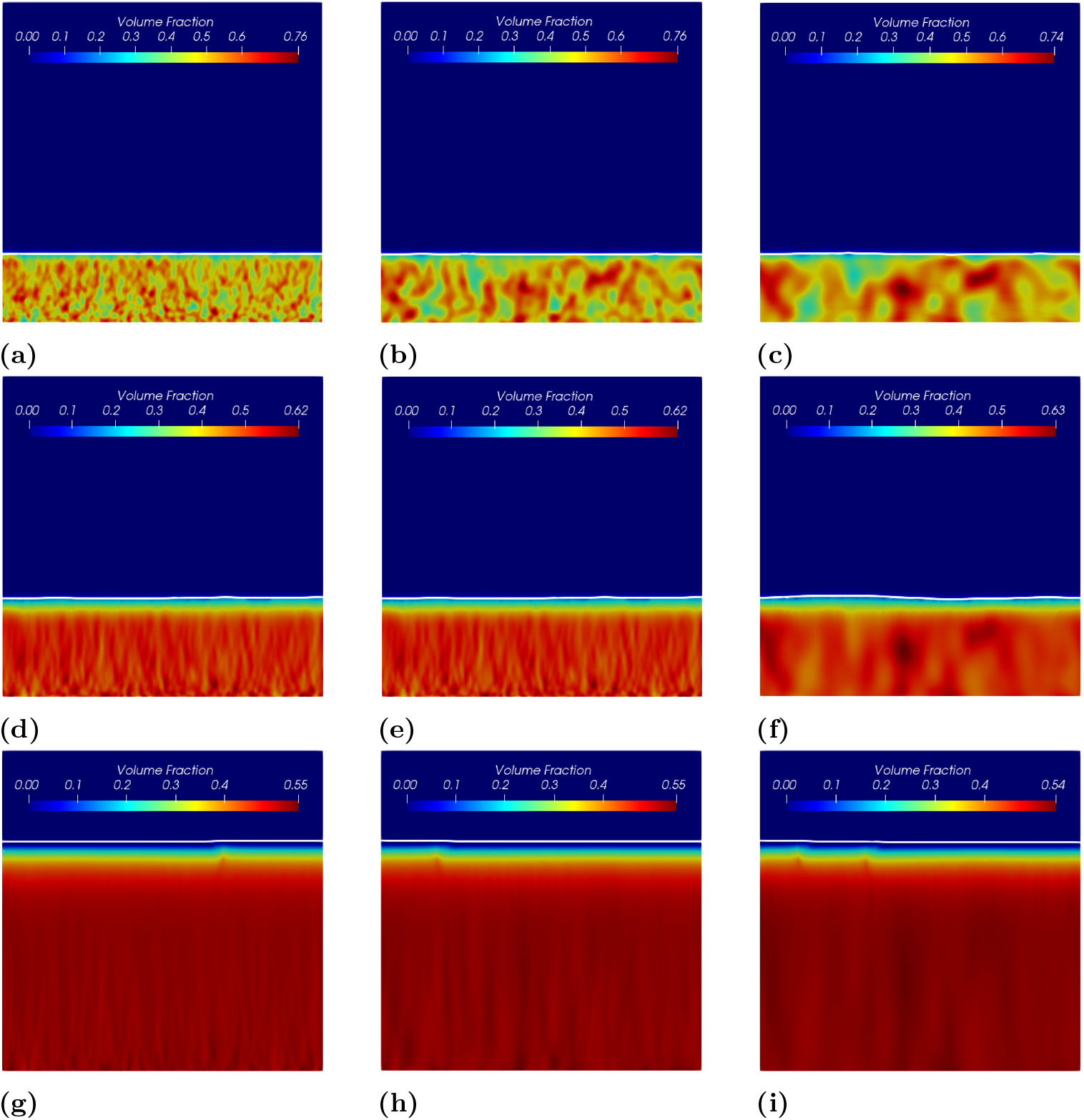
Volume fractions of *Veillonella* at different times corresponding to different initial distribution with various corrlation lengths. Figure from left to right refer to simulation results corresponding to different initial correlation lengths of 2Δ*x*, 4Δ*x* and 8Δ*x*. Figure from top to bottom refer to simulation results at different times of *t* = 0.1 day (*T* = 0.1), *t* = 0.5 day (*T* = 0.5) and *t* = 1.5 day (*T* = 1.5).

**Figure 13.**
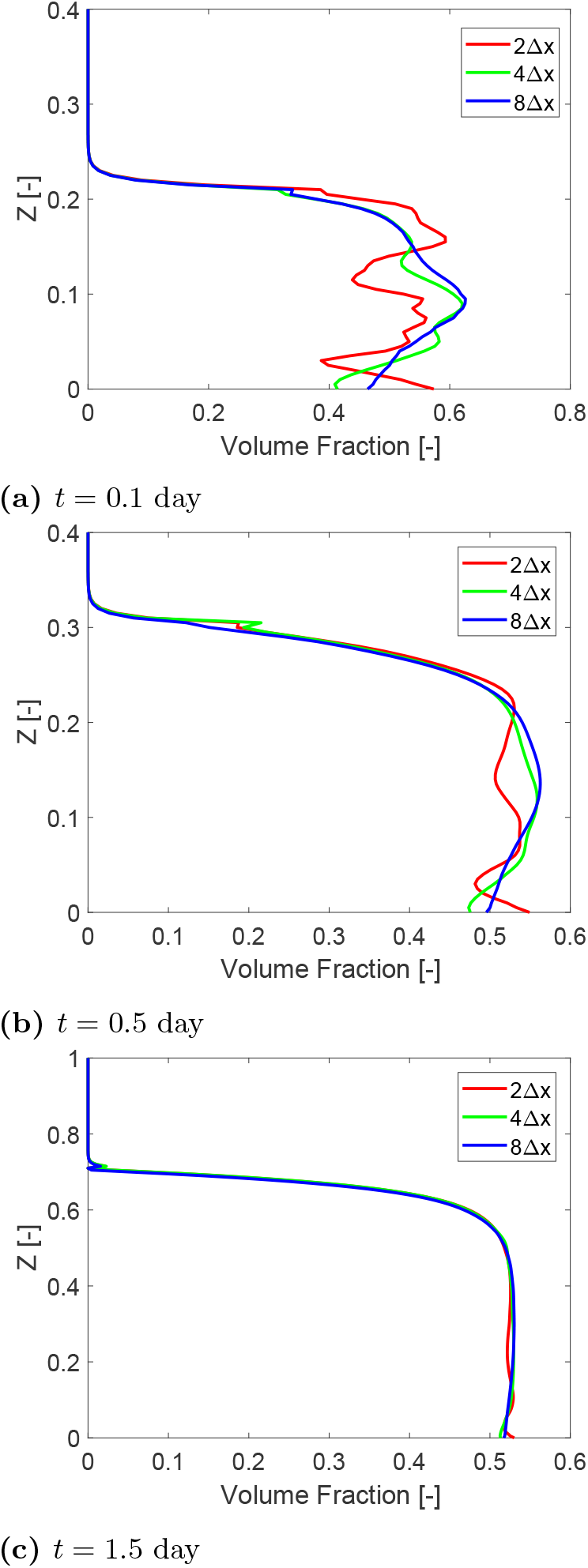
Volume fraction profiles of *Veillonella* at *X* = 0.5 at *t* = 0.1 day (*T* = 0.1; early time), *t* = 0.5 day (*T* = 0.5; middle time) and *t* = 1.5 day (*T* = 1.5; late time).

The evolution of *M*_diff_ corresponding to these three scenarios is illustrated in Figure 14. The results demonstrate that the numerical solutions are homogenized in all these three cases as soon as the simulation starts. The case corresponding to a smaller correlation length requires less time to reach the state of a given value of *M*_diff_ at the early time. In other words, the system homogenizes faster when the correlation length is smaller. However, as mentioned earlier, more fluctuated vertical biomass profiles are observed corresponding to the scenarios with larger correlation length. Moreover, *M*_diff_ increases at a later time (as shown in Figure 14). This is due to even though the initial biomass in the homogeneous case is the same as these three heterogeneous cases, the distribution of the lactic acid is different from case to case which results in different biomass production in the heterogeneous cases than in the homogeneous case. Such a difference may accumulate over time and leads to an increase in the measured mass difference *M*_diff_ at late time. In a word, the presented heterogeneous system will not converge to the homogeneous system where the biomass is initially evenly mixed.

**Figure 14.**
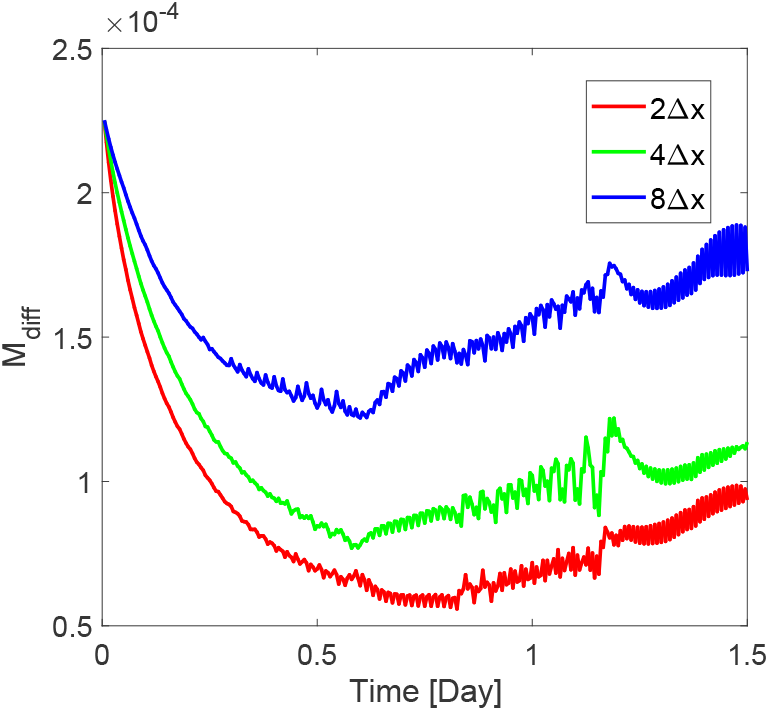
Homogenization times of three scenarios corresponding to initial biomass distribution with different correlation lengths 2Δ*x*, 4Δ*x* and 8Δ*x*.

### 3.3 Initial biomass distribution with patch-shape *Veillonella*

One could also argue that a random initial distribution of biomass is not realistic. It is more often found in mature biofilm systems that the *Veillonella* group bacteria are embedded in *Streptococcus* group bacteria (Chalmers et al., 2008; Kommerein et al., 2017). Therefore, we carry out studies with two different initial biomass distribution scenarios as shown in Figure 15. The irregular geometry of the *Veillonella* clusters, numbered from 1 to 4 (from left to right), are generated by using an implicit superquadric function which is defined by an implicit equation with random parameters (Wellmann and Wriggers, 2012) and is rotated by a random angle. In case a), the initial biofilm-fluid interface is flat, while a non-flat initial interface is assigned in case b).

**Figure 15.**
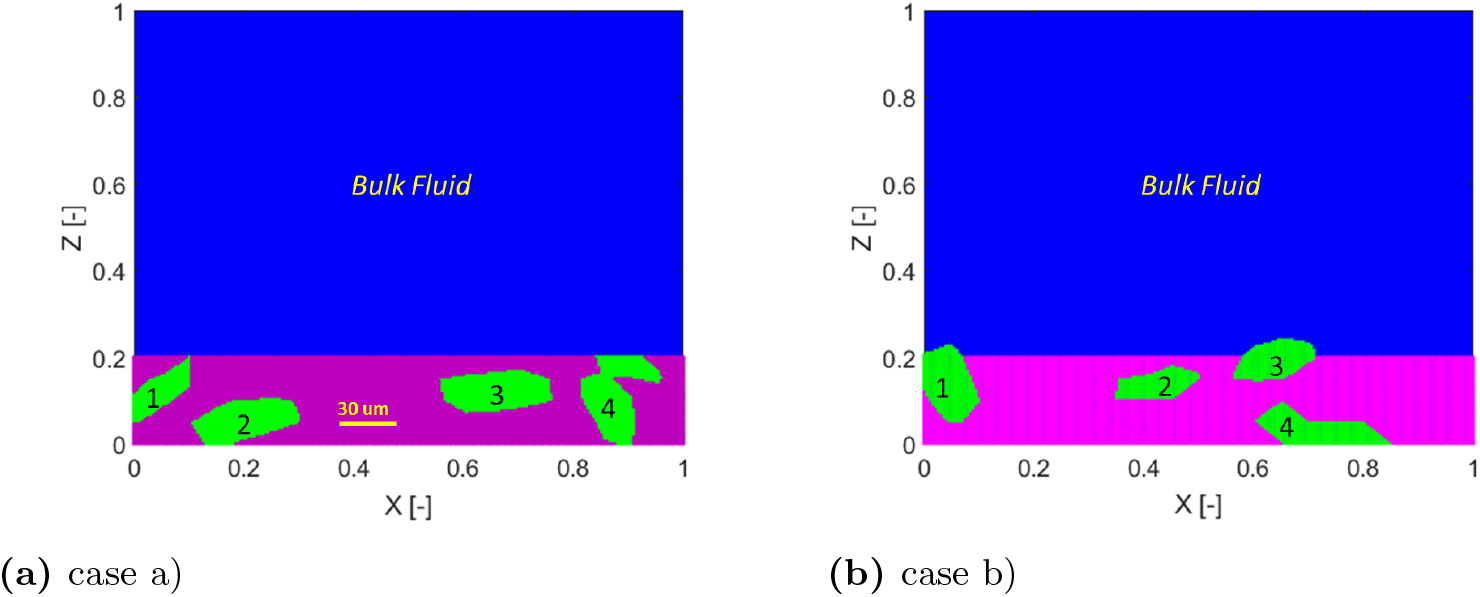
Two scenarios of initial biomass distribution (blue:bulk fluid; purple: *S. gordonii;* green: *Veillonella).*

Simulation results of these two cases with different initial biomass distributions are shown in Figure 16 and Figure 17. The simulation results demonstrate that the locally homogenization occurs at a late time (at 1.0 day). Some *Veillonella* patches are still observable even at *T* = 1.0. One may expect that the patches will be homogenized if we run the simulation long enough. Also, a non-flat biofilm-fluid interface is generated during the growth even if the interface is flat initially (as shown in Figure 16). This is due to the heterogeneous reactions, which result in a heterogeneous biofilm growth velocity field. With the homogenization of the solution, the biofilm interface gets flatted again

**Figure 16.**
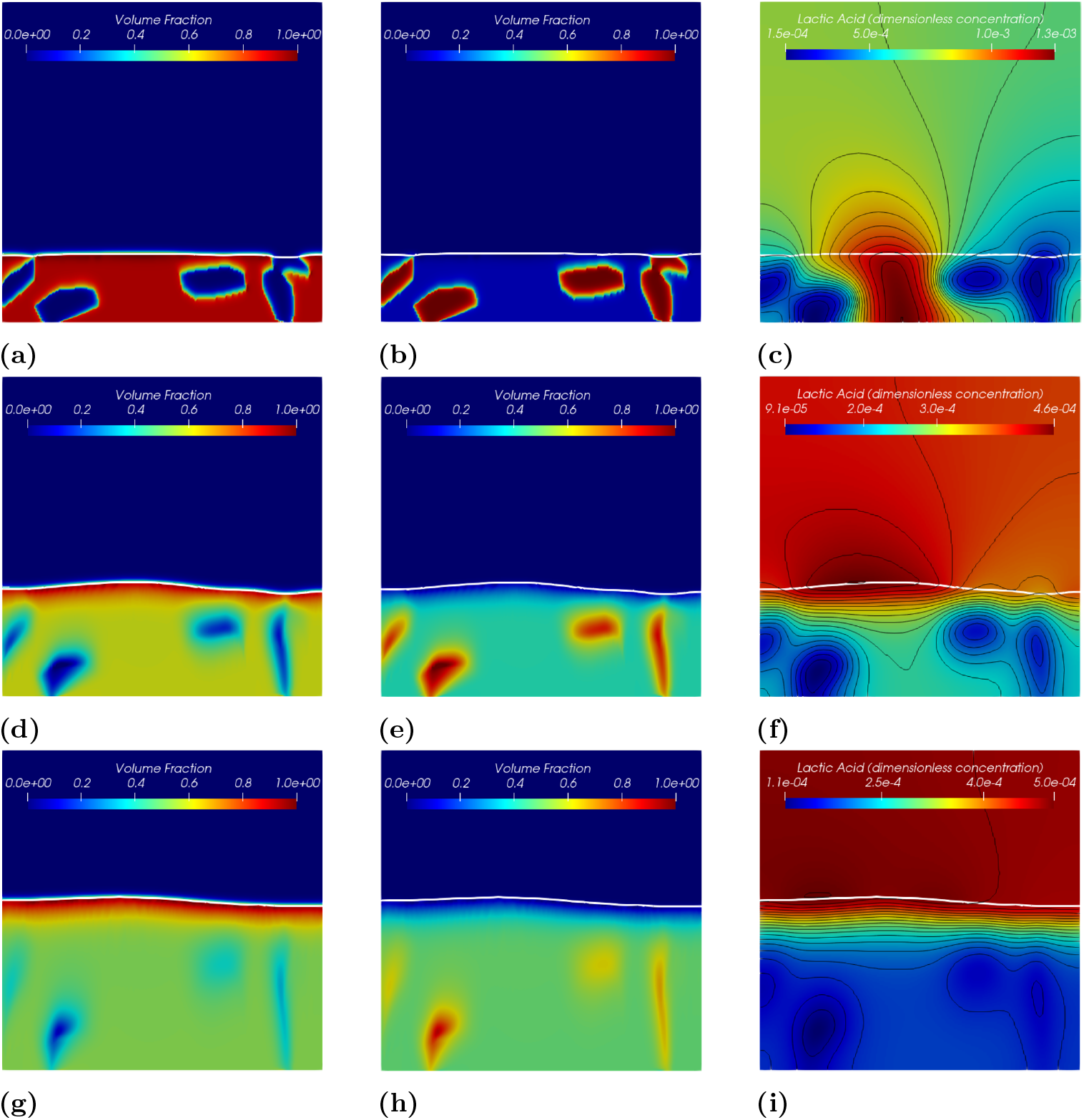
Simulation results of the symbiotic biofilm system at different times (corresponding to the case in Figure 15a). The white curve denotes the biofilm-fluid interface. From top to bottom are at *t* = 0.05 day (*T* = 0.05), *t* = 0.5 day (*T* = 0.5) and *t* = 1.0 day (*T* = 1.0). Left column: Volume fraction distributions of *S. gordonii;* Middle colume: Volume fraction distributions of *Veillonella;* Right column: Dimensionless concentration of lactic acid (black curves denote isolines of *S*_2_).

Contours of the volume fraction of *Veillonella ϑ*_2_ = 0.6 of each case at *t* = 0 and at *t* = 1.0 day are plotted in Figure 18. In Figure 18a, the second *Veillonella* cluster (numbered in Figure 15) seems to be rotated anticlockwise. However, the third cluster moves up together with the growth of the biofilm. The forth one is simply stretched and shrunken. In case b), as shown in Figure 18b, we observe an interesting feature of the numerical solution that the third and the forth *Veillonella* clusters cannot be captured by the contour line *d*_2_ = 0.6 at 1.0 day. This demonstrates that those two clusters are homogenized faster than the other two clusters. The simulation results suggest that the homogenization time varies spatially which might be induced by the heterogeneous reactions.

**Figure 17.**
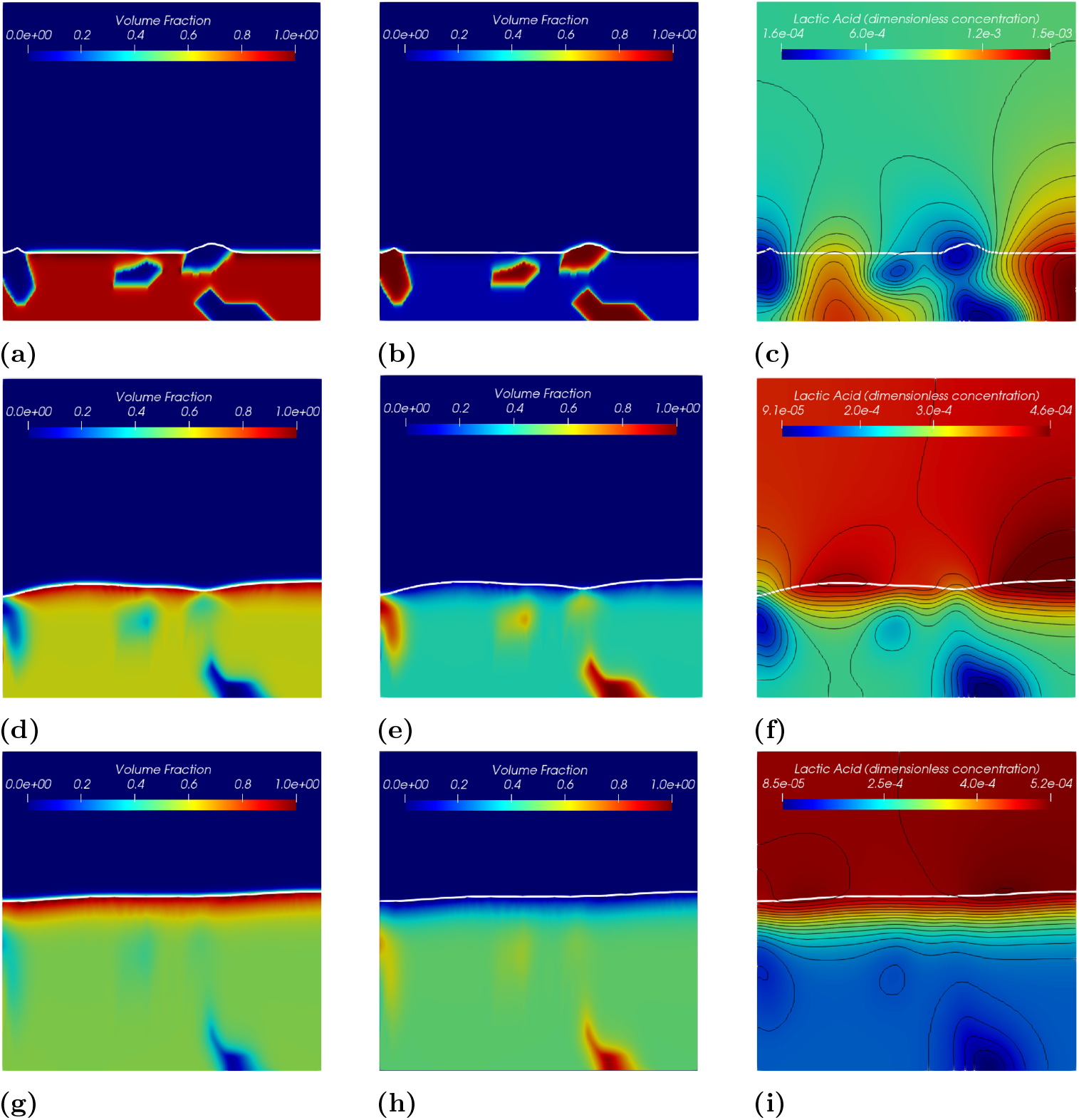
Simulation results of the symbiotic biofilm system at different times (corresponding to the case in Figure 15b). The white curve denotes the biofilm-fluid interface. From top to bottom are at *t* = 0.05 day (*T* = 0.05), *t* = 0.5 day (*T* = 0.5) and *t* = 1.0 day (*T* = 1.0). Left column: Volume fraction distributions of *S. gordonii;* Middle colume: Volume fraction distributions of *Veillonella;* Right column: Dimensionless concentration of lactic acid (black curves denote isolines of *S*_2_).

**Figure 18.**
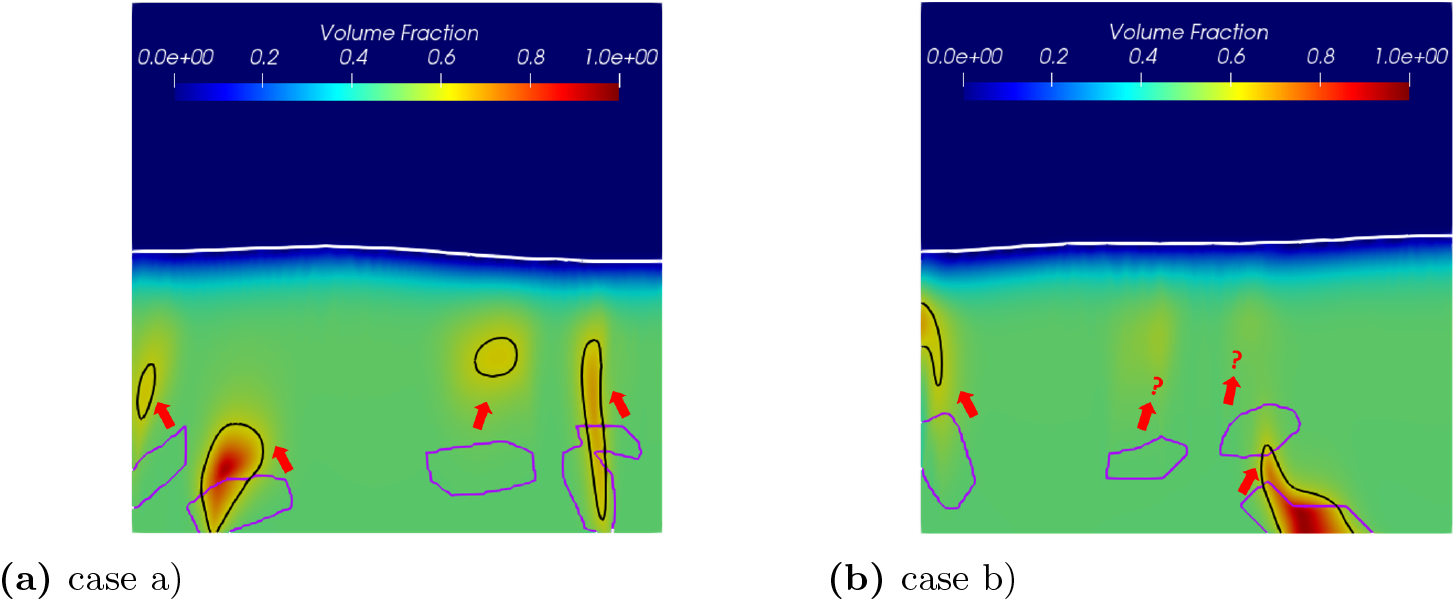
Movement of *Veillonella* clusters over time. The purper curve and the black curve denote contour lines of volume fraction of *Veillonella d*_2_ = 0.6 at *t* = 0 day (*T* = 0) and *t* =1.0 day (*T* = 1.0) respectively.

### 3.4 The role of the symbiotic reactions

Following the discussions in the previous section, the presented reaction model may be one of the main reasons that cause local biomass homogenization. To understand the role of the reaction terms in the mathematical model, we compare the simulation results presented in Figure 18b to the results by using a model without symbiotic reactions. The same model setup, as the case shown in Figure 18b, has been applied in this study. We simplify the symbiotic model (SM) presented in section 2 by applying:

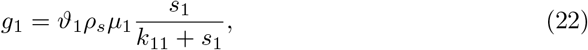

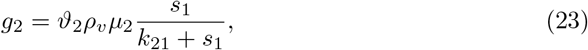

and

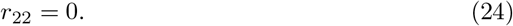

The model reduces to a competitive model (CM) in which the growth of both species of bacteria is limited only by the nutrient (saliva). In this case, the lactic acid does not play a role in biofilm growth, and it accumulates in space. One would expect that by applying CM, the biofilm grows faster than using SM due to the absent limitation of the lactic acid. Meanwhile, the species which has a larger maximum growth rate becomes dominant during the growth. As shown in Figure 19, the simulation results of the *Veillonella* sp. by using CM meet the above expectations. The *Veillonella* sp. becomes dominating during the growth and wins the competition. The simulation results of the CM (as shown in Figure 19) do not homogenized as in the SM (see Figure 18b). This demonstrates that the symbiotic interaction is one of the main reasons that lead to the homogenized solutions.

**Figure 19.**
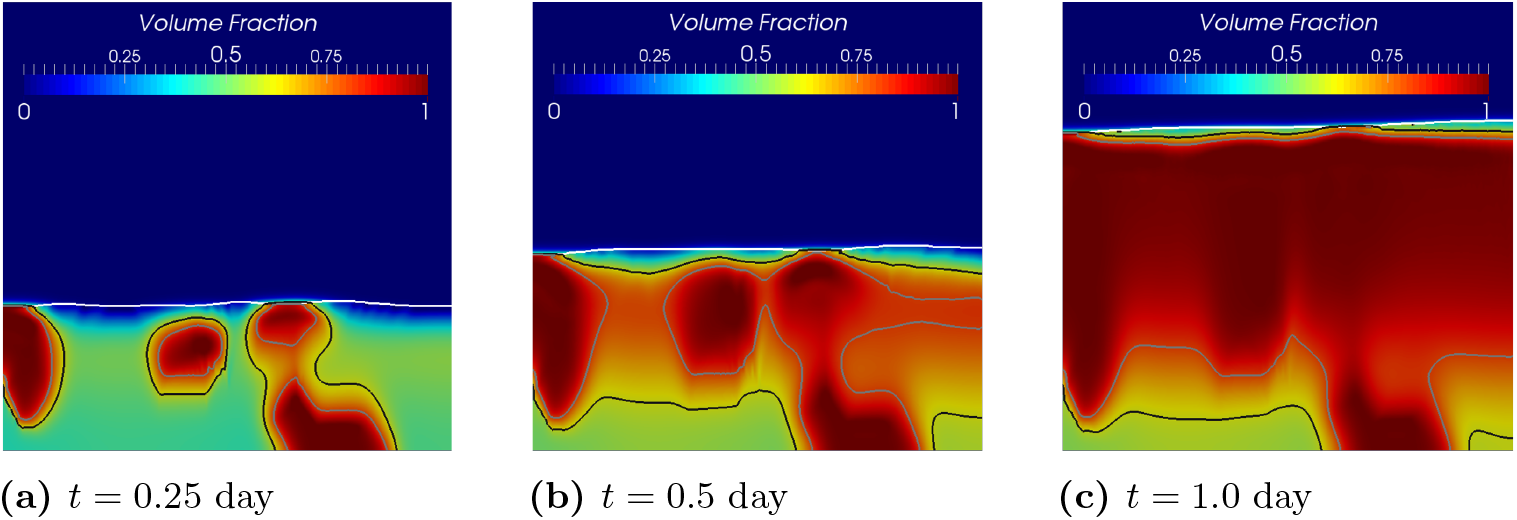
Simulation results of volume fractions of *Veillonella* sp. at different times by using the competitive model. The white curve denotes the biofilm-fluid interface. The black and grey curves denote contour lines of volume fraction of *Veillonella ϑ*_2_ = 0.6 and *ϑ*_2_ = 0.8 respectively.

## 4 Summary and discussion

### 4.1 Summary

We have presented a new mathematical model for modeling the growth of symbiotic biofilms in this paper. An intermediate product, namely the lactic acid produced by the *Streptococcus* sp., has been explicitly modeled. We found that the volume fractions of different species of bacteria homogenize at a later time. To understand the solution behaviors, we studied how the initial biomass distribution influences the homogenization process. Random distributions of the biomass volume fractions with different correlation lengths were taken as initial conditions. We compared the simulation results of the case where the bacteria were initially evenly mixed. As would be expected, the biomass homogenized faster associating with an initial state with a smaller correlation length. However, the random distribution cases do not converge to the same concentration distribution as the evenly mixed one, although the same initial mass and morphology of the biofilm surface (flat surface) have been assigned.

For a better understanding of the role of the initial distributions, we investigated scenarios with patch-shape biomass. The simulation results demonstrate that the local growth field can lead to changes in the morphology of the biofilm surface and different bacteria patches’ movements. Moreover, we also observe various homogenization speeds that are induced by the heterogeneity of the symbiotic reactions. To understand the role of the reactions in the homogenization process, we simplified the model by switching off the symbiotic reactions, resulting in a competitive model. The competitive model’s numerical solution does not get homogenized, which demonstrates that the symbiotic reaction is one of the main reasons for spatial homogenizations.

### 4.2 Discussion

It is well-known in microbiology that bacterial biofilms’ biological behaviors are a lot more complicated than a mere competition of “food” in nature. Many efforts have been put in the past decades to understand how different bacteria species interact in a biofilm. The symbiotic relationship between two species of bacteria is one of the most interesting phenomena in microbiology systems.

Mathematical modeling has been proved to be a powerful tool for understanding biofilm’s biological behaviors in the past years (Mattei et al., 2017). Tens or even hundreds of mathematical models have been developed for modeling the bacterial biofilm systems since the 1980s (Rittmann and McCarty, 1980). Nowadays, biofilm models are getting more comprehensive than earlier ones due to a deeper understanding of the biological mechanisms. Based on the previous pioneer continuum biofilm models (Alpkvist and Klapper, 2007; Klapper and Dockery, 2002), we studied a dual-species biofilm system’s symbiotic behavior by using mathematical modeling.

To the best of the authors’ knowledge, this study provides the first comprehensive investigation of the dynamics of a biofilm’s symbiotic reactions using a continuum mathematical model. We modeled the biofilm growth as an advective movement of a reactive potential flow in this study. The model has the advantage of capturing a sharp biofilm interface without introducing additional parameters; however, it suffers limitations on modeling merge of different colonies (see discussion in section 2). To overcome this drawback, one can improve the model by considering the fluid phase explicitly and thus lead to the mixture models (Zhang et al., 2008a). Alternatively, one can model the biofilm growth as a diffusion process (Ghasemi et al., 2018; Rahman et al., 2015), and apply the reaction models presented in this paper directly.

Instead of modeling the colonies’ merge, we are more interested in the reaction model developed for the symbiotic biofilm systems. Even though the simulation results demonstrate that the model can represent the species of bacteria’s symbiotic behaviors, we found that the reactive model can lead to homogenized solutions. The presented model may fail to predict the biological behavior of the symbiotic biofilms after a certain time when the numerical solution becomes over-homogenized. Therefore, it is necessary to investigate what factors influence the homogenization speed in the model. It turned out that the speed of the homogenization depends on the initial biomass distribution.

Individual bacteria are never resolved in a continuum biofilm model. This might be another potential reason which leads to the over-mixing of the biomass. However, the discretized element based biofilm models can also result in considerable internal mixing, especially for multi-species problems (Tang and Valocchi, 2013). It would be interesting to compare the mixing behavior occurred in different continuum and discretized element based biofilm models in future studies. As a matter of fact, there are still many open questions on modeling multi-species biofilm problems. A third possible reason that results in the unwanted homogenized solution is missing important bio-processes in the model. There could be specific quorum sensing (QS) mechanisms that prevent the mixing process in nature. Egland *et al.* presented experimental evidence of signaling between *S. gordonii* and *V. atypica* (Egland et al., 2004). The diffusive short distance signaling induced by the quorum sensing (QS) might be one of the key reasons that lead to the “corn cob” pattern of the biofilm system. Mathematical modeling the QS in biofilm systems has already attracted much attention in recent years (Emerenini et al., 2015; Ghasemi et al., 2018; Zhao and Wang, 2017). However, many more detailed QS mechanisms are still unclear (Egland et al., 2004) and may result in many uncertainties in the model.

## A Dimensioless form of governing equations

**Mass balance for saliva**

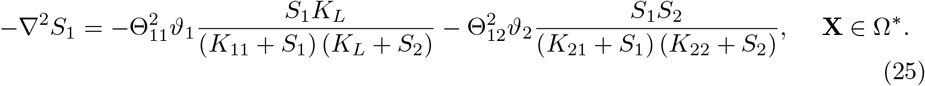

**Mass balance for lactic acid**

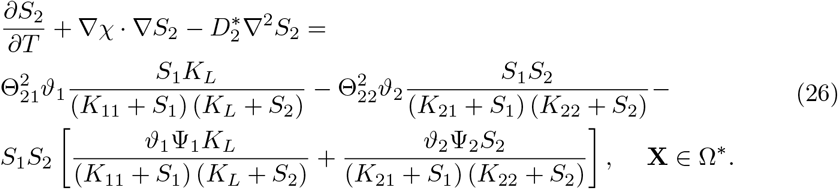

**Mass balance for** *S. gordonii*

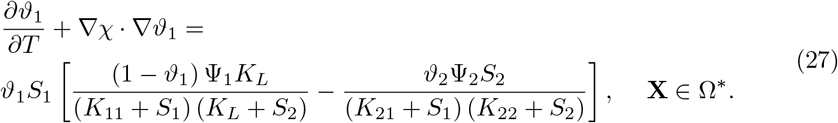

**Mass balance for** *Veillonella*

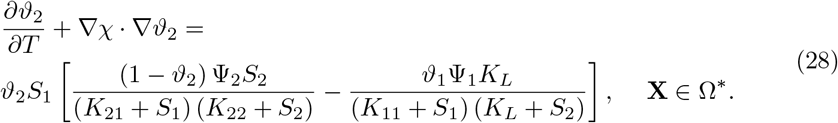

**Potential equation**

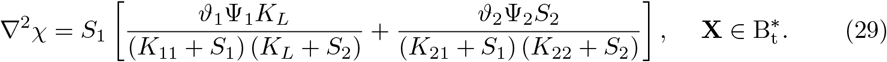

## B Numerical methods

Two types of PDEs, namely the elliptic equations (25) and (29) and the time dependent advection-diffusion-reaction (ADR) equations (27) and (28) (or advection-reaction equations) are involved in the current biofilm model. The mass balance equations of the lactic acid and the biomass (S. *gordonii* and *Veillonella)* are solved simultaneously in this study. The Time-discontinuous Galerkin (TDG) method (Hughes and Hulbert, 1988; Hulbert, 1992) is applied for solving the time dependent PDEs. The finite increment calculus (FIC) method (Oñate, 1998; Oñate et al., 2007) is used to stabilize the numerical solutions. We would like to refer to (Feng et al., 2017; Sapotnick and Nackenhorst, 2012) for more detailed interpretation of the TDG-FIC method on solving the time dependent ADR equations. In this study, 4-node bi-linear iso-parametric elements are applied for spatial discretization of the mass balance equations of the saliva, lactic acid and biomass. 8-node second order iso-parametric elements are used for solving the potential equation. The free boundary (moving biofilm-fluid interface) at each time step is captured by using an iso-line of 80% of total biomass volume fraction. Detailed information on the numerical schemes involved in the numerical strategy will be presented in the following part. As a remark, only one and two-dimensional problems are studied in this paper even through the mathematical model also can be applied for three-dimensional problems.

### B.1 Finite element approximation of nonlinear Poisson equation with *C*^0^ elements

Equations (25) and (29) are Poisson equations. In this study, they are solved by using conventional Galerkin finite element method with *C*^0^ type of elements. Both of these two equations can be written generally as

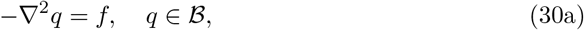

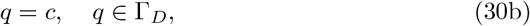

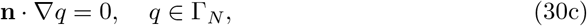

where *q* represents the primary variables as *S*_1_ or *χ* in the governing equations. *f* denotes a function as 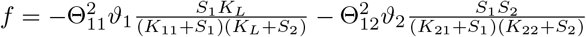 for equation (25) and 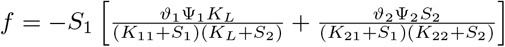 for equation (29). 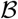 refers to the computational domain with the corresponding Dirichlet and Neuman boundaries Γ_*D*_ and Γ_*N*_. **n** is the norm vector on the Neuman boundary and *c* is a constant value corresponding to the Dirichlet boundary. The weak form of equation (30) can be written as

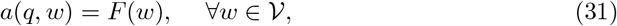

where,

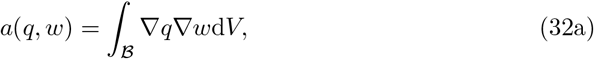

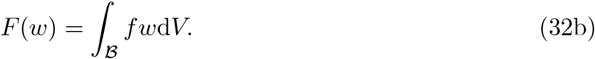

*w* is an arbitrary test function in space 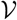 which is a subspace of the Hilbert space 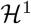. As a remark, the Newton-Raphson method (Wriggers, 2008) is used when the equation is nonlinear.

### B.2 TDG-FIC approximation of time dependent advection-diffusion-reaction equation

Equations (26)–(28) are time-dependent PDEs. Therefore, they are solved simultaneously by using the TDG-FIC method. Those three equations can be written as a set of time dependent advection-diffusion-reaction partial differential equations as

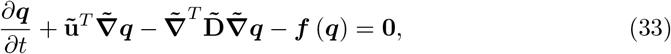

where ***q*** = (*S*_2_, *ϑ*_1_, *ϑ*_2_)^*T*^ denotes the unknown vector. Detailed information on the matrix form of each term in equation (33) can be found in C.

Applying discontinuous Galerkin method for time discretization and standard Galerkin discretization in space yields a time-space weak form of equation (33) over a time-space domain 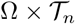 (see Figure 2 in (Feng et al., 2019))

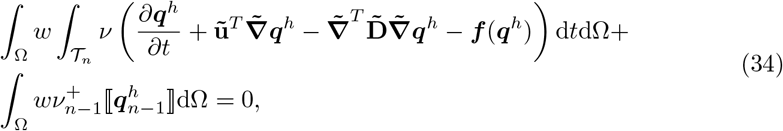

where Ω and 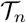 denote the spatial and temporal computational domains, respectively. ***q***^*h*^ denotes the time-space approximation of the variable vector. Since the nodal value at a time point is discontinuous, a jump value of 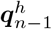 at *t*_*n*-1_, which is notated as 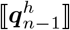, is introduced as

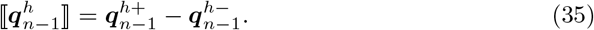

Due to the non-linearity of the reaction terms ***f*** (***q***^*h*^), the Newton-Raphson method (Wriggers, 2008) is applied and equation (34) is linearized by introducing an incremental process with ***q***^*h*,0^ = **0** as

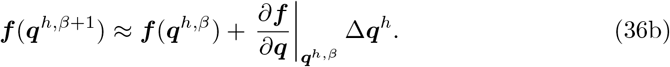

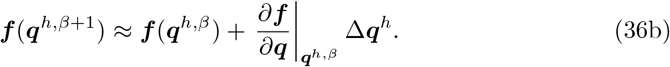

Substituting (36) into (34) yields the linearized time-space weak form

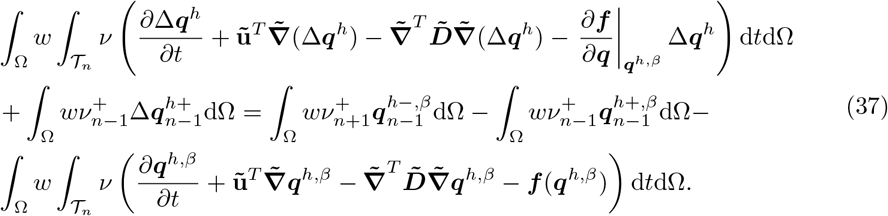

The unknown field of ***q***^*h*^ is interpolated over the time-space space by

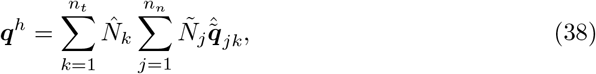

where *n_t_* and *n_n_* refer to temporal and spatial total degrees of freedoms respectively. 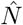 and 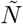 are the temporal and spatial shape functions and 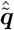 denotes the time-space nodal variable vector.

Discretization of equation (37) in space and time by adopting equation (38) within single time-space element 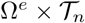 yields

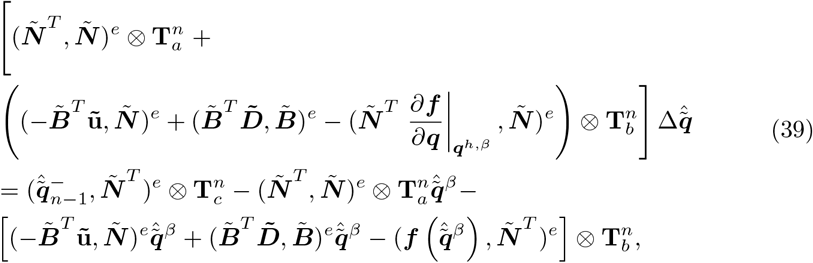

where

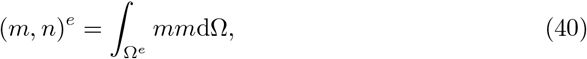

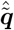 refers to the time-space approximation of the nodal variables. **T**_*a*_, **T**_*b*_, **T**_*c*_ are time matrixes with only temporal shape function 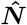 involved

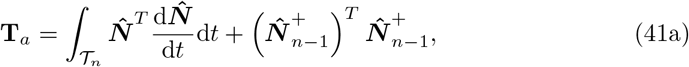

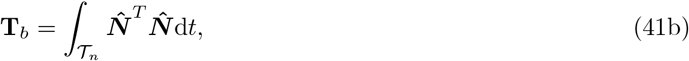

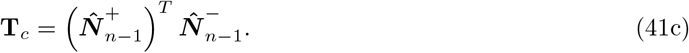

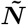 is extended spatial shape function depending on the spatial element type (see C for detailed matrix form of 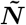).

The diffusion term in (33) only affects the mass balance equation of the lactic acid and does not appear in the mass balance equations of biomass. Therefore, the system of the mass balance equations in this model is still hyperbolic dominated. It is well known that solving hyperbolic PDEs with the conventional Galerkin method suffers from convective instability (Donea and Huerta, 2003). One way to relieve such a problem is by adding an additional artificial diffusion (balancing diffusion) term into the system. This yields the so-called “stabilization methods” (Codina, 1998; Franca et al., 1992; Lian et al., 2016) which have been developed for Galerkin methods over the past decades. In this paper, one of the widely used stabilization methods, namely the finite increment calculus (FIC) method, which introduces a nonlinear balancing artificial diffusion term in (39) is adopted. Thus, the diffusion coefficient matrix 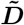 is replaced by

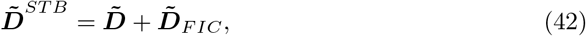

where

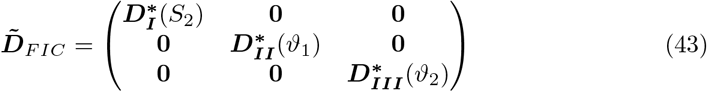

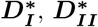 and 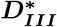 are 2 × 2 diagonal matrices depending on the variables *S*_2_, *ϑ*_1_ and *ϑ*_2_ respectively as in (43). Detailed calculation procedures of these ***D***\* matrices for multi-dimensional problems can be found in (Feng et al., 2017). As a remark, 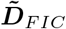 depends on the solution of each spatial element at the current time step. Moreover, the artificial balancing diffusion is introduced along the directions of the principle curvatures of the solution and thus it must be calculated in a local coordinate system and then transferred to the global Cartesian coordinate system afterwards. Therefore, the FIC stabilized algorithm becomes nonlinear and iterations at each time step are required even if a linear problem is considered.

## C Matrices in the advection-reaction equation

For two-dimensional problems, the advection and diffusion coefficients matrices read

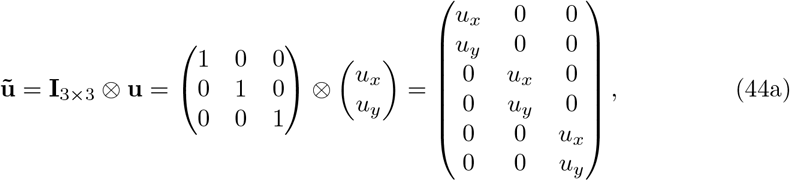

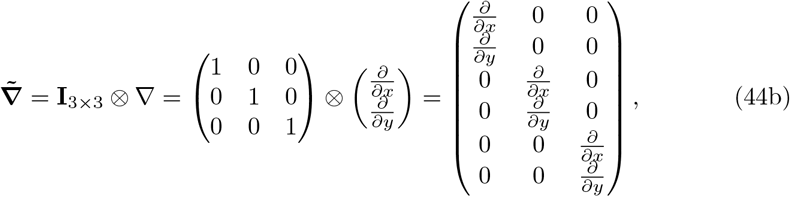

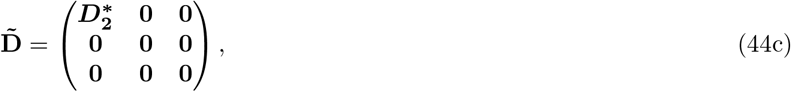

where *u_x_* and *u_y_* are the advection velocity components along *x* and *y* coordinates respectively. **I** denotes an identity matrix and ⊗ is the Kronecker product. **I** denotes an identical matrix and ⊗ is the Kronecker product. 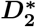 denotes a 2 × 2 diagonal matrix with 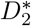 placed at the main diagonal and **0** denotes a 2 × 2 null matrix. Furthermore, *f* (***q***) reads

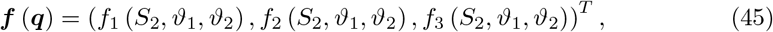

where *f*_1_, *f*_2_ and *f*_3_ denote the reaction terms of equations (26)–(28) respectively.

For four-node 2D bi-linear element, the shape function is extended to

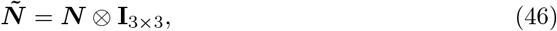

where ***N*** = [*N*_1_, *N*_2_, *N*_3_, *N*_4_] is the shape function of the four-node 2D bi-linear element. Similarly,

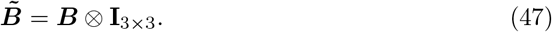

## Acknowledgements

The authors would like to thank Dr. Henryke Rath, Dr. Nadine Kommerein and Mrs. Natascha Brandhorst for many helpful discussions. Dianlei Feng would also like to thank the German Research Council (DFG) under grant No. 426819984.

